# The gut microbiome has sexually dimorphic effects on bone tissue energy metabolism and multiscale bone quality in C57BL/6J mice

**DOI:** 10.1101/2022.11.07.515521

**Authors:** G. Vahidi, M. Moody, H. Welhaven, L. Davidson, S. T. Walk, S. Martin, R. K. June, C. M. Heveran

## Abstract

The gut microbiome impacts bone mass, implying a disruption to bone homeostasis, yet significant uncertainty remains regarding the impacts of the gut microbiome on remodeling bone cells. The gut microbiome is thought to be essential for normal biomineralization, but the specific consequences of the absent gut microbiome on tissue mineralization and multiscale bone quality are not determined. We hypothesized that bone homeostasis and tissue-scale metabolism, tissue mineralization, and whole-bone biomechanics are altered in germ-free (GF) C57BL/6J mice. Further, because many characteristics of the gut microbiome are sexually dimorphic, we hypothesized that the gut microbiome would show important sex differences with regards to its impact on bone quality. Differences between GF and conventional mouse bone extended from bone tissue metabolism to whole bone biomechanics. Cortical bone tissue from male mice had a greater signature of amino acid metabolism whereas female cells had a greater signature of lipid metabolism. These sex differences were also present in GF mice and were indeed even more stark. GF increased cortical femur bone formation for both sexes and decreased bone resorption and osteoclast density only in females. GF similarly increased cortical femur tissue mineralization and altered collagen structure for both sexes but led to greater gains in distal femur trabecular microarchitecture for males. Whole femur strength was similarly increased with GF for both sexes, but males had a greater increase in modulus. GF did not impact fracture toughness for either sex. The altered bone quality with GF is multifactorial and is likely contributed to by differences in tissue-scale composition as well as lower cortical porosity. Together, these data demonstrate that the gut microbiome influences bone cells and multiscale bone quality, but that the specific relationships that underlie these effects to bone are different for females and males.

## 1. Introduction

The mammalian gut microbiome is composed of trillions of microbial cells and is responsible for the production of a diverse set of molecules^(1)^. Evidence suggests that the composition of the gut microbiome can drive sex-dependent differences in host phenotype and disease^(2–6)^. Moreover, gut microbiota composition and inflammatory phenotypes associated with the gut flora are themselves sexually dimorphic^(2, 7–11)^. The repertoire of gut microbial antigens and metabolites can influence bone mass through their impacts on nutrient transport, immune system regulation, and translocation of bacterial products into the systematic circulation and bones ^(9, 12–17)^. However, whether the microbiome has sexually dimorphic effects on bone cells, bone tissue metabolism, and multiscale bone quality is still uncertain, which may presently mask important interactions between the gut and the skeleton.

Evaluations of hindlimbs from germ-free (GF) mice show that the gut microbiome is essential for normal bone homeostasis. In several studies, female GF C57BL/6 mice had increased bone mass, trabecular microstructure, and cortical geometry compared to conventionally raised female mice^(10, 18–21)^ (**Table 1**). Though the increased bone mass of GF mice implies that the activities of osteoblasts and osteoclasts are dysregulated, the specific impacts of the gut microbiome on the abundance and activities of each of these cells are not clear. Because the GF immune system fails to mature^(9, 18, 22–24)^, osteoclast maturation would be expected to be decreased. However, while Sjorgen and coauthors reported a decrease in osteoclast abundance at the femur of 9 weeks-old GF mice^(18)^, Li *et al* reported no change in osteoclast abundance at the femur of 12 weeks-old GF mice^(19)^. Similarly, Novince *et al* reported higher expression of osteoblast related genes and proteins such as Runx2, Col12a, and osteocalcin in marrow cell cultures from femur of 12 weeks-old GF mice^(25)^, but Yan and coauthors reported lower expression of Runx2 in epiphyseal bone from 13 weeks-old GF mice^(26)^ (**Table 1**). Therefore, whether and to what extent osteoblast and osteoclast abundance and activity in GF mice differ from those of conventional mice remains unclear.

**Table 1.**
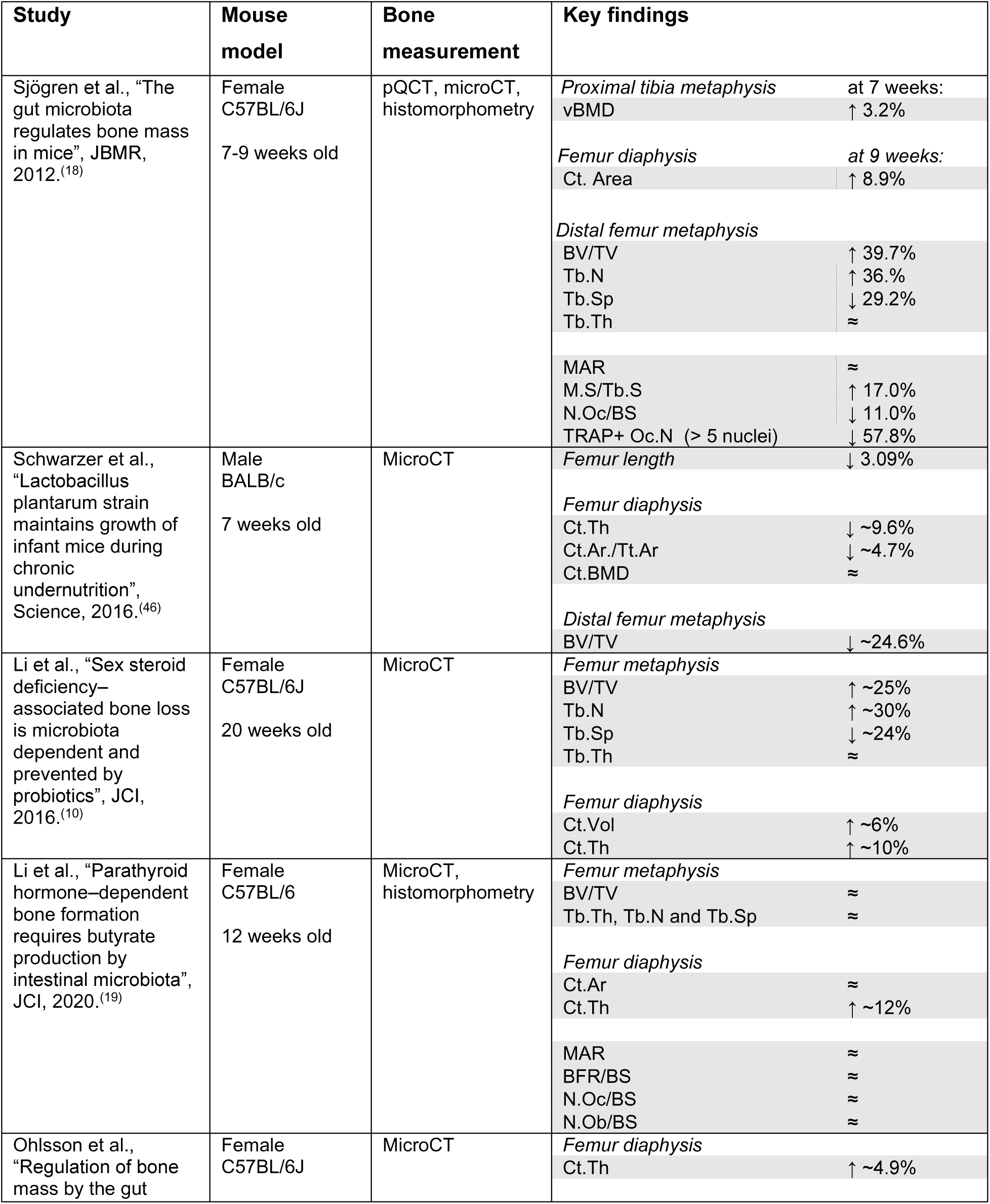
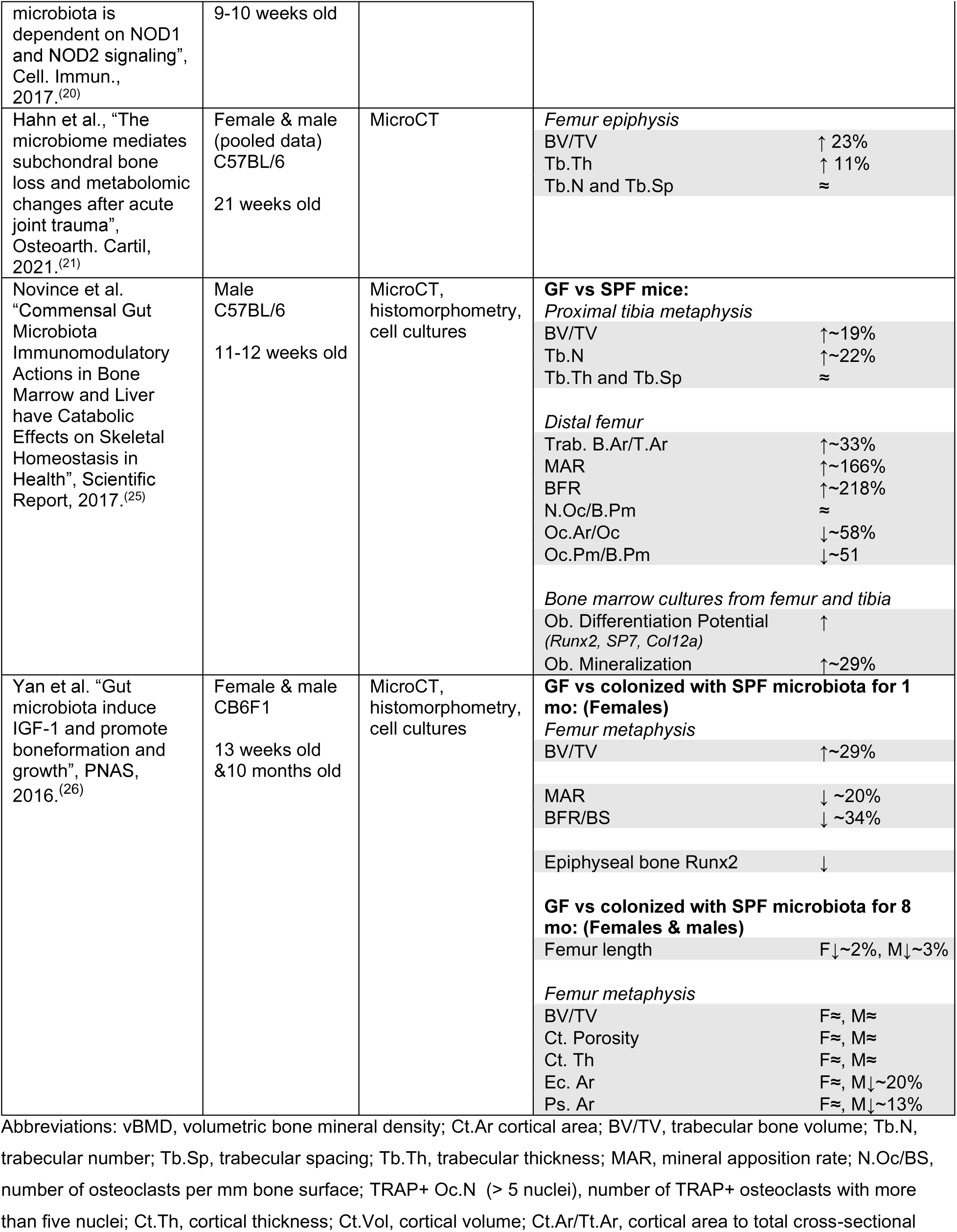
Literature review of the effects of germ-free (GF) status on hindlimb bone quality. Arrows directions are in reference to the effect of GF versus conventionally raised mice.

Even less is known regarding osteocyte abundance and function in the GF state. For example, GF mice lack bacteria-driven vitamin K biosynthesis^(9)^, which has been shown to play an important role in osteoblast to osteocyte transition^(27–31)^. Therefore, it is possible that GF mice have less osteocytes. Currently, it is unknown whether osteocyte abundance, signaling, or perilacunar remodeling are disrupted in GF mice and whether these changes may also be dependent on sex. This particular knowledge gap is important because osteocytes are essential for indirectly and directly contributing to the regulation of bone mass and bone quality over the lifespan^(32, 33)^, and often have sexually dimorphic characteristics^(34–36)^.

Whether bone material properties in addition to bone mass and microarchitecture are altered in the GF model is not understood (**Table 1**). The absent gut microbiome must necessarily eliminate the production of gut microbe-derived forms of vitamin K (menaquinones MK5-MK13, otherwise known as vitamin K_2_)^(37)^. Vitamin K_2_ directly impacts bone mineralization through carboxylating osteocalcin, the most abundant non-collagenous protein^(27, 38–41)^. Some work also indicates that changes to osteocalcin mineralization can deleteriously affect fracture toughness^(27, 38, 42, 43)^. However, it has not been evaluated if the GF mouse has altered tissue-scale biomineral quantity or mineral maturity, and whether these changes are translated into differences in whole bone fracture resistance.

If the microbiome is important in bone cell physiology, it is reasonable that at least some of the pathways involved depend on microbial metabolites either used directly by host cells for their own metabolism or indirectly as metabolic regulators as is the case for other non-bone tissues^(44)^. Thus, there is a premise for interrogating whether bone tissue metabolism is also regulated by the microbiome. Studying the metabolism of bone tissue provides a snapshot of cellular level bioenergetics, which aids in the interpretation in differences in bone remodeling activity. We recently found that cortical bone metabolic pathways were sexually dimorphic in 20- week-old C57BL/6J mice^(45)^. Female mice had greater levels of lipid metabolism while male mice had higher levels of amino acid metabolism. Stronger bones in both females and males had higher tryptophan and purine metabolism^(45)^. Assessing bone tissue metabolism for GF and conventional mice of both sexes provides new insight into the connections between the microbiome, bone cell health, and bone quality.

In the present study, we hypothesized that the absence of the gut microbiome in C57BL/6J mice increases bone mass and strength and that these differences extend to increased tissue-scale mineralization, altered bone cell remodeling, and dysregulated bone tissue metabolism. We further hypothesized that these effects of gut microbiome on the skeleton interact with sex.

## 2. Materials and Methods

### 2.1 Animal model

All animal procedures were approved by Montana State University’s Institutional Animal Care and Use Committee. Female and male C57BL/6J mice (female; n = 6, male; n = 7) were reared in standard cages inside a hermetically sealed isolator with HEPA-filtered airflow and maintained on sterile (autoclaved) water and food (LabDiet® 5013, Land O’Lakes) ad libitum. GF status was confirmed using standard cultivation and molecular biology techniques^(47)^. Briefly, liquid ‘bug’ traps comprised of a mixture of drinking water and food were left open to the air inside of isolators and observed daily for signs of microbial growth (i.e., turbidity). Stool samples from mice were monitored prior to and throughout experiments for signs of growth on rich media under anaerobic and regular atmosphere conditions (Mueller–Hinton broth and agar plates). Bulk DNA was also extracted from stool samples (DNeasy PowerSoil Pro DNA isolation kit, Qiagen, Hilden, Germany) to serve as a template for PCR using primers targeting the bacterial 16S rRNA encoding gene. GF status was confirmed through lack of growth and amplification by PCR. Age- and sex-matched conventionally-raised C57BL/6J mice (female; n = 10, male; n = 10) were also used. Conventional mice were housed in cages of 3-5 mice and fed a standard chow diet ad libitum (LabDiet® 5053, Land O’Lakes). Alizarin label (30 mg/kg; SIGMA: A3882-1G) was administered, via intraperitoneal injection, 3 days before euthanasia. Animals were euthanized by cervical dislocation at age 20-21 weeks.

### 2.2 Quantitative Reverse Transcription Polymerase Chain Reaction (qRT-PCR)

Marrow-flushed left tibiae were pulverized in liquid nitrogen and homogenized in Trizol (Life Technologies). Total RNA was isolated using a Qiagen RNeasy Mini Kit (Qiagen) according to the manufacturer’s protocol. RNA was reverse transcribed into cDNA using a High-Capacity cDNA RT kit (ThermFisher Scientific). qRT-PCR gene expression analyses were conducted on an Applied Biosystems QuantStudio 5 platform using PR1MA qMax Gold SYBR Green Master Mix. Gene expression for RankL (receptor activator of nuclear factor–kappa B Ligand), MMP2 (matrix metalloproteinases), MMP13 (Matrix metalloproteinase-13), MMP14 (Matrix metalloproteinase-14), OPG (osteoprotegerin), ACP (TRAP, Tartrate-resistant acid phosphatase), and CTSK (cathepsin K) was determined using the following primer sequences (**Table 2**). Target gene expression was normalized to 18S, and relative quantification was determined (ΔΔCt method). The RankL/OPG ratio was determined using ΔCt calculations.

**Table 2.**
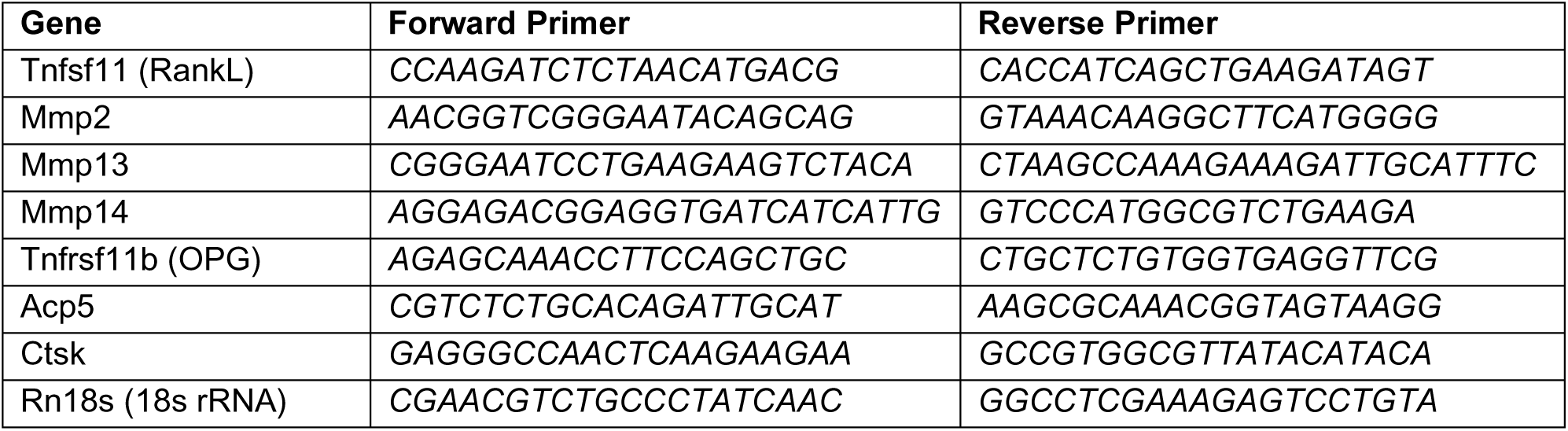
Primer sequencing used for PCR analysis.

### 2.3 Histology

For histological analysis, right tibiae were decalcified with EDTA disodium salt dihydrate, dehydrated in a graded ethanol series, embedded in paraffin, and serially sliced into 5-micron thick horizontal cortical sections. Sections from each sample were stained with terminal deoxynucleotidyl transferase dUTP Nick End Labelling (TUNEL) and tartrate resistant acid phosphatase (TRAP). Two cortical sections were analyzed per sample for both TUNEL and TRAP stains. Histological slides were imaged using a Nikon E-800 microscope (Nikon, Melville, NY) with 4x and 10x objectives. All image analysis was performed using Fiji ImageJ software. Total lacunae and empty lacunae were measured from 10x TUNEL-stained sections. Number of osteoclasts and pink stained lacunae were obtained from TRAP-stained sections. Images taken using the 4x objective were used to determine total cortical area (TUNEL) and endocortical perimeter (TRAP).

Bone marrow adiposity was measured as previously described^(45)^, using hematoxylin and eosin (H&E) staining on longitudinally cut, 5 µm-thick sections of tibia. Sections were imaged, and bone marrow adiposity was quantified through manually segmentation and a custom MATLAB code to obtain measurements including mean adipocyte area (mm2), marrow cavity area (mm2), adipocyte count, and adipocyte number density (number of adipocytes per marrow cavity area).

### 2.4 Serum chemistry analysis

Serum collected via cardiac puncture at euthanasia was assessed for biomarkers of bone remodeling (P1NP and CTX1). Serum P1NP was measured using a Mouse procollagen 1 N-terminal peptide (P1NP) ELISA kit (MBS703389, My BioSource) and CTX1 was measured using a Mouse Cross linked C terminal Telopeptide of type 1 collagen (CTX1) ELISA kit (MBS722404, My BioSource) according to the manufacturer’s protocols.

### 2.5 Quantitative histomorphometry

An upright confocal laser scanning microscope (Leica SP3, Heildelberg GmbH, Mannheim, Germany) was used to visualize the alizarin fluorochrome labeled periosteal and endocortical perimeters of the embedded cortical femur (sample preparation procedure is described later). Imaging was performed with the following parameters: 5x objective, laser wavelength excitation 633 nm (emission length 580-645), 600 Hz speed with a 1024 × 1024 resolution, pinhole set at 1 Airy unit, and laser intensity set at 50% of the full power. The gain and offset were set to the best label visibility and minimum noise per sample. ImageJ software was used for image processing. Confocal images were converted to a maximum contrast to visualize reliable labeled bones and achieve a consistent thresholding. Then, the perimeter of alizarin labeled (AL.Pm) bone for endocortical and periosteal surfaces was measured. Total endocortical and periosteal bone perimeters (Tt.Pm) were also calculated from confocal images. Percentage mineralizing surface (MS/BS=(AL.Pm/Tt.Pm)×100) was reported for endocortical and (Ec.MS/BS) periosteal surfaces (Ps.MS/BS) for each group. Some animals that did not receive labels were excluded from this analysis.

### 2.6 Trabecular microarchitecture and cortical geometry

A high-resolution desktop micro-tomographic imaging system (μCT40, Scanco Medical AG) was used to assess trabecular microstructure and cortical geometry of femurs. Left femurs were harvested and fresh frozen at -20°C in phosphate-buffered saline (PBS) soaked gauze before microCT analysis. Scans were acquired using a 10 μm^3^ isotropic voxel size, 70 kVP, 114 μA, and 200 ms integration time. Scans were subjected to Gaussian filtration and segmentation. Image acquisition and analysis protocols adhered to JBMR guidelines^(48)^. Trabecular microarchitecture was evaluated at the femoral distal metaphysis in a region beginning 200 μm superior to the top of the distal growth plate and extending 1500 μm proximally. Endocortical region of the bone was manually contoured to identify the trabeculae. Trabeculae was segmented from soft tissue with a 375 mgHA/cm^3^ threshold. Using the Scanco Trabecular Bone Morphometry Evaluation Script, following architectural parameters were measured: bone volume fraction (BV/TV, %), trabecular bone mineral density (BMD, mgHA/cm^3^), connectivity density (Conn.D, 1/mm^3^), structural model index (SMI), trabecular bone surface to bone volume ratio (BS/BV, mm^2^/mm^3^), trabecular thickness (Tb.Th, mm), trabecular number (Tb.N, mm^-1^), and trabecular separation (Tb.Sp, mm).

Cortical geometry was evaluated at the femoral mid-diaphysis in 50 transverse micorCT slices (500 μm long region) in a region including entire outermost edge of the cortex. Cortical bone was segmented with a fixed threshold of 700 mgHA/cm^3^. Following cortical parameters were measured; cortical bone area (Ct.Ar, mm^2^), medullary area (Ma.Ar, mm^2^), total cross-sectional area (bone + medullary area) (Tt.Ar, mm^2^), cortical tissue mineral density (Ct.TMD, mgHA/cm^3^), cortical thickness (Ct.Th, mm), minimum moment of inertia (I_min_, mm^4^), polar moment of inertia (pMOI, mm^4^), and the maximum radius perpendicular to the I_min_ direction (C_min_, mm).

### 2.7 Whole-bone mechanical and material properties

The left femurs were assessed for flexural material properties using three-point bending (1 kN load cell, Instron 5543, Norwood, MA). The test was performed on PBS-hydrated femurs to failure at a rate of 5 mm/min on a custom fixture with 8 mm span. Femurs were positioned such that the posterior surface in tension. Load-displacement data, and I_min_ and C_min_ values from micorCT were used based on standard equations for the mouse femur^(49)^ to calculate material and mechanical properties of the femur such as modulus, maximum load, maximum strength, yield strength, post yield displacement, work to fracture, and toughness (i.e., area under stress-strain curve until first failure).

Notched fracture toughness was evaluated for right femurs, consistent with our description in Welhaven *et al*^(45)^. A custom device (**Supp. Figure 1**) was used to notch the posterior surface of mid-shaft femurs to a target notch depth was 1/3 of the anterior-posterior width^(50)^. Bone hydration was maintained using PBS. Notched femurs were then tested in three-point bending (1 kN load cell, Instron 5543) at a rate of 0.001 mm/s on a custom fixture with 8 mm span until failure^(50)^. Femurs were tested with the posterior surface in tension. Following the test, distal femurs were cleaned of marrow near the fracture surface and air dried overnight. Fracture surfaces were imaged using field emission scanning electron microscopy (FESEM, Zeiss SUPRA 55VP) in variable pressure mode (VPSE, 20 Pa, 15 kV). A custom MATLAB code was used to assess cortical geometry and initial notch half angle. Fracture toughness values (K_c-max_ and K_c-yield_) were calculated using the maximum load and yield load methods^(50)^ (**Equation 1**). The notch geometry satisfied the thick-wall cylinder criteria proposed by Ritchie *et al*^(50)^.

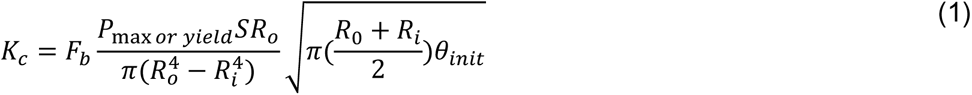

**Figure 1.**
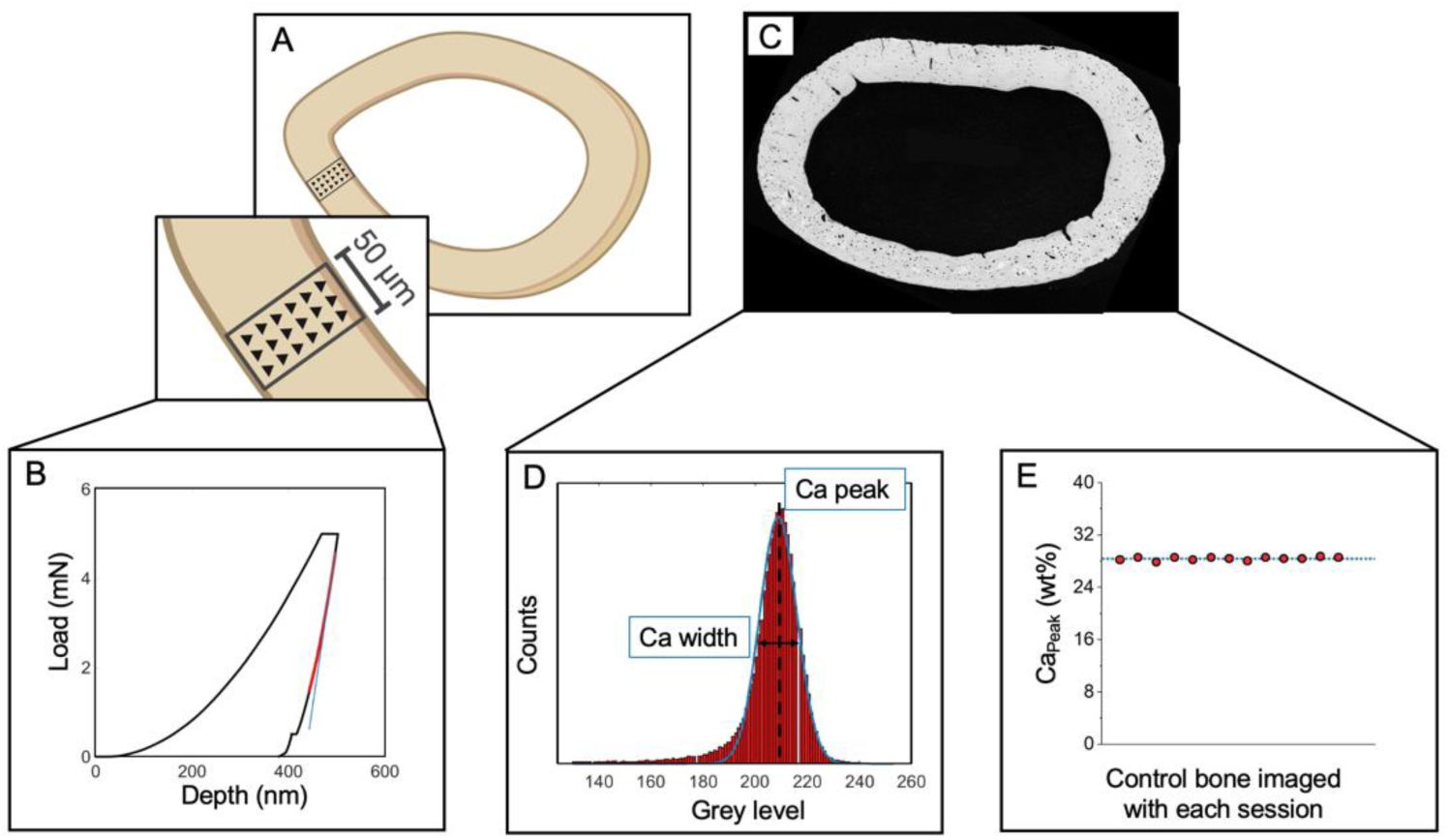
Tissue-scale characterization of the cortical femur mid-diaphysis. A) Tissue-scale modulus was assessed in maps spanning the cortical thickness in the posterior quadrant. B) Representative load-displacement curve from a nanoindentation test. C) Tissue mineralization was evaluated using quantitative backscattered electron microscopy. D) Ca_Peak_ Peak and Ca_Width_ were calculated from histograms of each whole femur cross-section. E) In addition to the use of reference standards, one control bone sample was evaluated with each imaging session. There was 0.85% variation in the Ca_Peak_ for the control bone measured across imaging sessions.

Where F_b_ is the geometry constant for thick-walled cylinders, P_max_ or P_yield_ are the max load or yield load, R_0_ and R_i_ are mean outer and inner radii, S is the span of loading (8 mm), and θ_init_ is the initial notch half angle.

### 2.8 Microscale assessment of cortical femur tissue modulus

PMMA-embedded left femurs were used for assessment of bone tissue modulus. Following whole-bone mechanical testing, left femurs were histologically dehydrated in a graded ethanol series (EtOH 70-100%) and embedded in poly(methyl) methacrylate (PMMA). Embedded femurs were sectioned at the midshaft using a low-speed diamond saw (Isomet, Buehler, Lake Bluff, IL) and the cortical surfaces were polished for further analyses. The polishing procedure included 600 and 1200 grits of wet silicon carbide papers (Buehler, Lake Bluff, IL) followed by fine polishing with Rayon fine clothes (South Bay Technologies, San Clemente, CA) and a series of alumina suspensions (9, 5, 3, 1, 0.5, 0.3, and 0.05 μm). Between each step, samples were sonicated with tap water to remove the remaining alumina particles.

Nanoindentation (KLA Tencor iMicro, Milpitas, CA) was performed on the posterior quadrant of each femur using a Berkovich tip. The target load was 5 mN and was applied with a load function of 30 s load, 60 s hold to dissipate viscoelastic energy before unloading^(51)^, and 30 s unload. Each nanoindentation map included three columns of indents spanning the whole cortical thickness (15 μm spacing in x and y; **Figure 1A**). The mean and standard deviation of the nanoindentation modulus were calculated for each femur using the Oliver-Phar approach^(52)^. The 95^th^-45^th^ percentile of the unloading curve was fit with a 2^nd^-order polynomial. A tangent line to the beginning of this section was used to calculate the stiffness (*S,* the slope of the unloading curve evaluated at the maximum load, 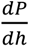, **Figure 1B**). The tip contact area (*A_c_*) was calculated as a function of the contact depth. The tip area was calibrated using fused silica (KLA Tencor, Milpitas, CA). The reduced modulus, *E_r_*, was calculated from *S* and *A_c_* as described in **Equation 2**.

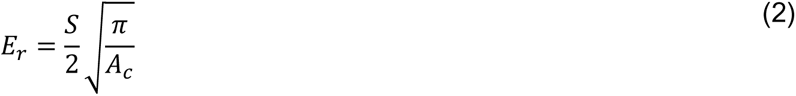

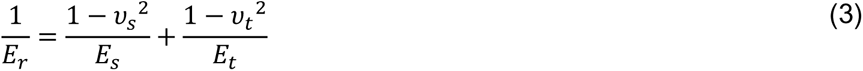

Nanoindentation modulus (*E_i_*) was then calculated from **Equations 3-4**, where the subscript *s* refers to the sample under study*. E_t_* and *ν_t_* are the known tip modulus (1140 GPa) and Poisson’s ratio (0.07), respectively. Since the sample’s Poisson’s ratio (*ν_s_*) is unknown, we report the indentation modulus *E_i_*, eliminating errors from an assumption for *ν_s_* (**Equation 4**).

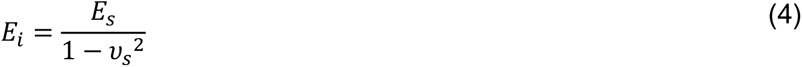

### 2.9 Microscale assessment of bone mineralization and porosity

Following nanoindentation, samples were coated with a thin layer of carbon for quantitative backscattered scanning electron microscopy (qBSE, Zeiss Supra 55VP field emission SEM, 20 kV, 60 μm aperture size, 100x magnification, and 9.1 mm working distance)^(53–55)^. A custom steel sample holder equipped with springs that pushes polished embedded bone samples against a flat steel plate was used to assure both flat sample surfaces and consistent working distances (**Supp. Figure 2**). BSE images of the cortical cross-sections were collected at 100x magnification (**Figure 1C**). Polished carbon and aluminum reference standards (Electron Microscopy Services) were mounted on the sample holder and imaged with bone samples with each imaging session. Images were processed by setting the mean grey levels of the aluminum and carbon calibration standards to 255 and 0, respectively^(56)^. A custom MATLAB code was used to convert the BSE images to corresponding calcium concentration, where each step in the greyscale corresponds with an increase of 0.1385 weight % calcium. Histograms of bone mineral density distribution with bin size of 1 grey level were generated for each calibrated image. From histograms, Ca_Peak_, the most frequent calcium concentration of the cortical surface (histogram peak), and Ca_Width_, heterogeneity of the Ca concentration within each sample (full-width at half-maximum of the histogram^(57)^ were calculated (**Figure 1D**). To assess variation between imaging sessions, we imaged one bone sample at each of the twelve imaging sessions and calculated the coefficient of variation (standard deviation/mean) in Ca_Peak_ measurement of this control bone. We observed 0.85% variability in Ca_Peak_ for this control bone between imaging sessions (**Figure 1E**)

**Figure 2.**
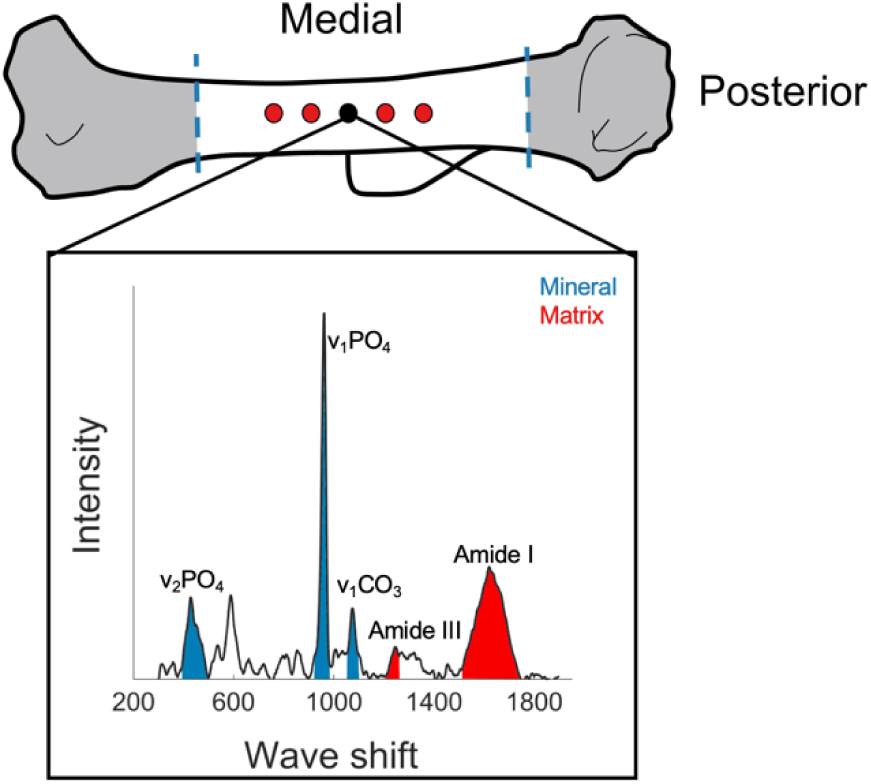
Raman spectroscopy of hydrated humeri. Spectra were collected from five points on the posterior side of humeri. First point (black-colored point) was located where the deltoid tuberosity connects to posterior side, and other points were located 50 μm apart spanning both directions.

Cortical porosity was assessed for each bone from a 400x image of the posterior cortical surface taken via secondary electron mode (SE2, Zeiss Supra 55VP, 20 kV, 30 μm aperture size, 9.1 mm working distance). A custom MATLAB code was used to calculate the total porosity (%) and pore number density (number of pores per area of interest, 1/mm^2^). Pores greater than 150 pixels^2^ were considered vasculature and pores smaller than this number were considered lacunae.

### 2.10 Microscale assessment of bone composition and collagen structure

Tissue composition and collagen properties were assessed using Raman spectroscopy (Horiba Laser Tweezer Raman Modified LabRam HR Evolution NIR) on hydrated right humeri. Humeri were thawed, cleaned, and flushed of marrow. For each sample 5 spectra were collected from the posterior side, located 50 μm apart. For each sample, the location of the deltoid tuberosity was used as a marker for the first spectra point (black point in **Figure 2**) and other points were spaced approximately 50 micrometers apart. Raman parameters were: 10x dry objective lens, 785 nm edge laser at 100% power, 300-1900 nm Raman spectra, 30 accumulations for 4 second acquisition time. Bones were maintained hydrated during the test using a sponge bed and tap water. Spectra data were baseline-corrected and analyzed using a custom MATLAB code. Mineral-to-matrix ratio (ν_2_PO_4_ / Amide ΙΙΙ), Carbonate substitution (ν_1_CO_3_ / ν_1_PO_4_, indicative of the extent of carbonate substitution into the mineral crystal lattice), crystallinity (full-width at half maximum of the ν_1_PO_4_ peak, FWHM [ν_1_PO_4_]^-1^), Amide I sub-band ratio at I_1670_/I_1690_ (indicative of enzymatic crosslinking in the secondary structure of collagen), and Amide I sub-band ratios at I_1670_/I_1610_ and I_1670_/I_1640_ (indicative of collagen helical status) were calculated^(58, 59)^ .

### 2.11 Evaluation of cortical bone metabolism

To investigate the metabolism of cortical bone, humeri-derived metabolites were subjected to liquid chromatography-mass spectrometry (LC-MS) and global metabolomic profiling were employed for the cortical bone of the humerus as previously reported^(45)^. Humeri ends were trimmed, and they were flushed of marrow with PBS to isolate cortical bone and then stored fresh frozen at -20°C in PBS-soaked gauze prior to analysis. Next, humeri were placed in liquid nitrogen for 2 hours and pulverized to optimize metabolite extraction. Pulverized bone was then precipitated with methanol:acetone, vortexed for 1 minute, and incubated at -20°C for four minutes. This process was repeated five times then samples were incubated overnight at -20°C to promote precipitation. The following day, samples were centrifuged, supernatant was dried down via vacuum concentration, and once dry, samples were suspended in acetonitrile:water.

Samples were analyzed using LC-MS (Agilent 6538 Q-TOF mass spectrometer) in positive mode (resolution: ∼20 ppm, adducts: H+, Na+) using a Cogent Diamond Hydride HILIC chromatography column, as previously described^(45, 60, 61)^. Agilent Masshunter Qualitative software, XCMS, MetaboAnalyst, and MATLAB were used for data analysis. Raw data were log transformed and auto-scaled (mean centered divided by standard deviation per variable) prior to analysis. Statistical analyses included hierarchical cluster analysis (HCA), principal component analysis (PCA), partial least squares-discriminant analysis (PLS-DA), volcano plot analysis, t-test, and fold change. MATLAB was utilized to examine differences in metabolite intensity across experimental groups. MetaboAnalyst’s Functional Analysis tool was used to identify biologically relevant pathways that are dysregulated between experimental groups (GF, sex, and GF-sex interactions).

### 2.12 Statistical analysis

Two-way ANOVA tested whether bone characterization outcomes depended on microbiome status (GF vs. conventional), sex (female vs. male), or their interaction (Minitab, v.20). Dependent variables were transformed, if necessary, such that all models satisfied assumptions of residual normality and homoscedasticity. Significance for main effects was set *a priori* to p < 0.05. Significant interactions between GF and sex were followed up with post-hoc tests. Family-wise type I error was maintained at 0.05 by using a Bonferroni correction. For most measures, GF vs conventional was compared within each sex (i.e., two comparisons, critical α: 0.05/2 = 0.025). Nanoindentation and Raman measurements were averaged per mouse such that one mean and one standard deviation for each measure per mouse were input into ANOVA models.

## 3. Results

### 3.1 Microbiome-sex interaction affects body weight but not femur length

GF and sex had an interactive effect (p = 0.001) on terminal body weights such that GF females were heavier than conventional females (+23.4%, 18.6%, p < 0.001) but weights were not different between GF and conventional males. Femur length was similar across groups (**Table 3**).

**Table 3.**
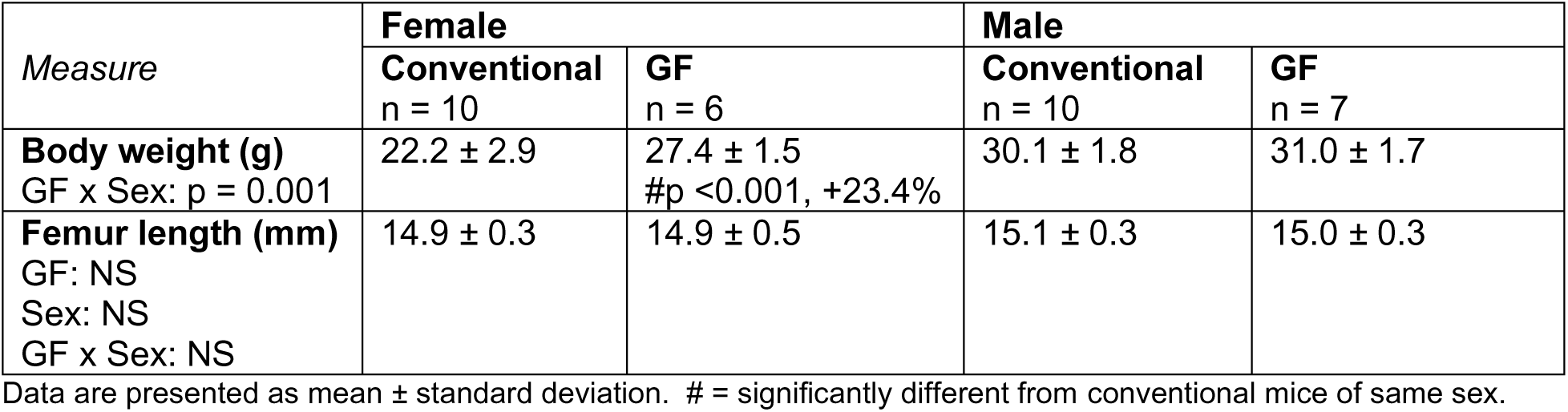
Terminal body weight and femur length.

| Measure | Female |  | Male |  |
| --- | --- | --- | --- | --- |
|  | Conventional<br>n = 10 | GF<br>n = 6 | Conventional<br>n = 10 | GF<br>n = 7 |
| **Body weight (g)**<br>GF x Sex: $p = 0.001$ | 22.2 ± 2.9 | 27.4 ± 1.5<br>#$p < 0.001$, +23.4% | 30.1 ± 1.8 | 31.0 ± 1.7 |
| **Femur length (mm)**<br>GF: NS<br>Sex: NS<br>GF x Sex: NS | 14.9 ± 0.3 | 14.9 ± 0.5 | 15.1 ± 0.3 | 15.0 ± 0.3 |

Data are presented as mean ± standard deviation. # = significantly different from conventional mice of same sex.

### 3.2 Microbiome and sex independently affect gene expression related to bone turnover

OPG expression from the marrow-flushed tibia was lower in GF mice compared to conventional mice (-51.6%, p = 0.032) (**Figure 3**). RankL expression was similar among groups. The RankL/OPG ratio was higher in GF mice compared to conventional mice (+127.6%, p = 0.033). We also assessed the expression of several genes involved in osteocyte perilacunar remodeling. MMP2 expression decreased with GF (-39.7%, p = 0.044) and the interaction between GF and sex on MMP2 expression was not significant (p = 0.07). MMP13 expression was similar among groups. MMP14 was expressed more in females compared to males (+64.0%, p = 0.03) but was unchanged with GF. CTSK expression was lower in females than males (+112.8%, p = 0.031) but not changed with GF. ACP5 did not differ with sex or GF. Gene expression data are reported in **Supp. Table 1**.

**Figure 3.**
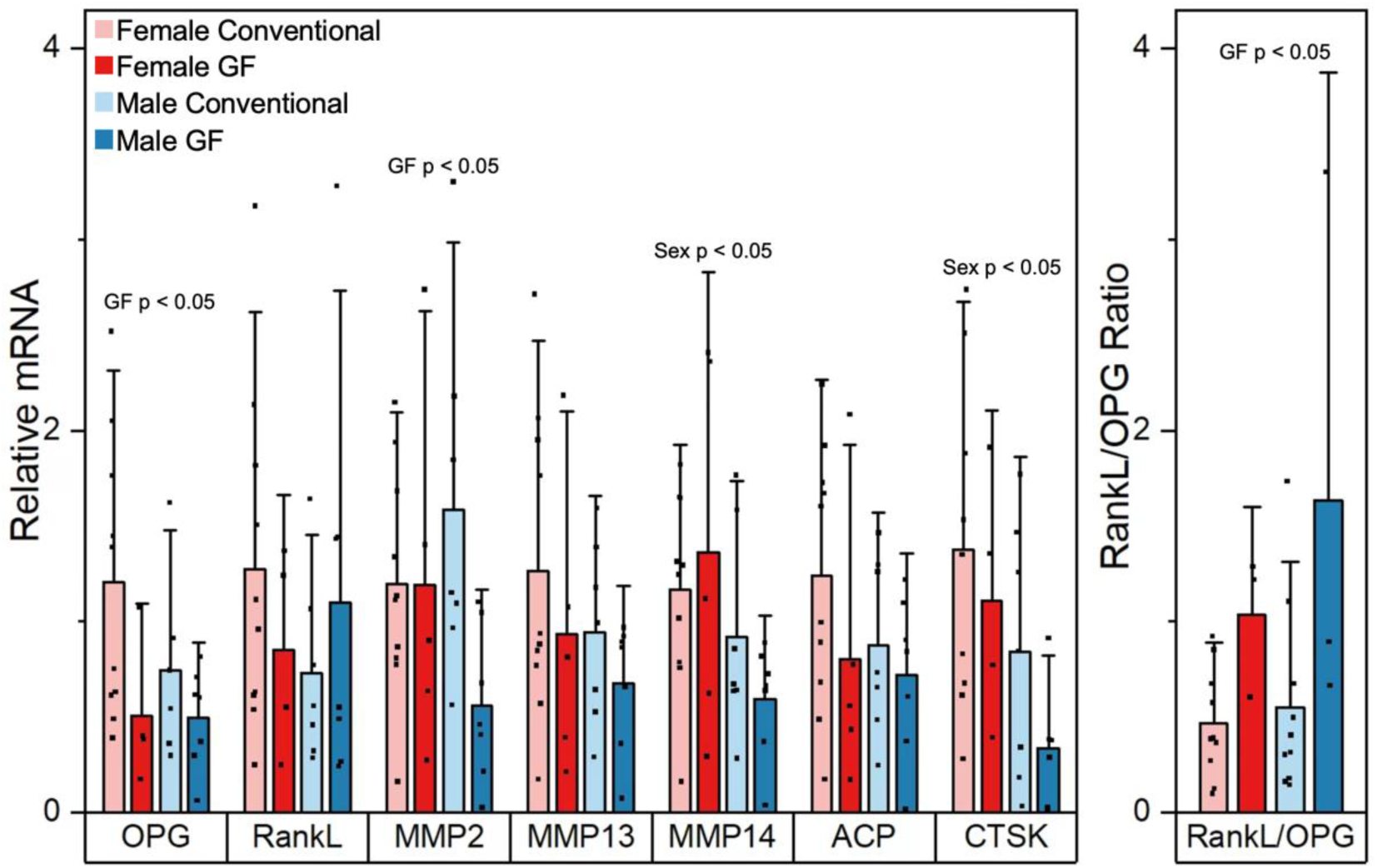
Effects of GF and sex on relative gene expression levels (fold changes) of OPG, RankL, MMP2, MMP13, MMP14, ACP and CTS, and on non-relative expression level of RankL/OPG ratio. Data are presented as means. Error bars indicate one standard deviation. P-values for significant main effects of GF or sex are shown above each gene. There were no interactions between sex and GF.

### 3.3 Microbiome-sex interaction affects osteoclast and osteocyte bone resorption

Sex and GF had an interactive effect on osteoclast number density, such that only GF females had reduced osteoclasts per endocortical perimeter compared to their conventional mice (-75.5%; p = 0.022, **Figure 4A**). Osteocyte perilacunar bone resorption, as estimated from TRAP-positive lacunae, was decreased in females and GF overall (-67%, p = 0.043; -155%, p = 0.021, respectively, **Figure 4B**). In contrast, lacunar number density and percent empty lacunae were not influenced by microbiome status or sex (**Figure 4C-D**).

**Figure 4.**
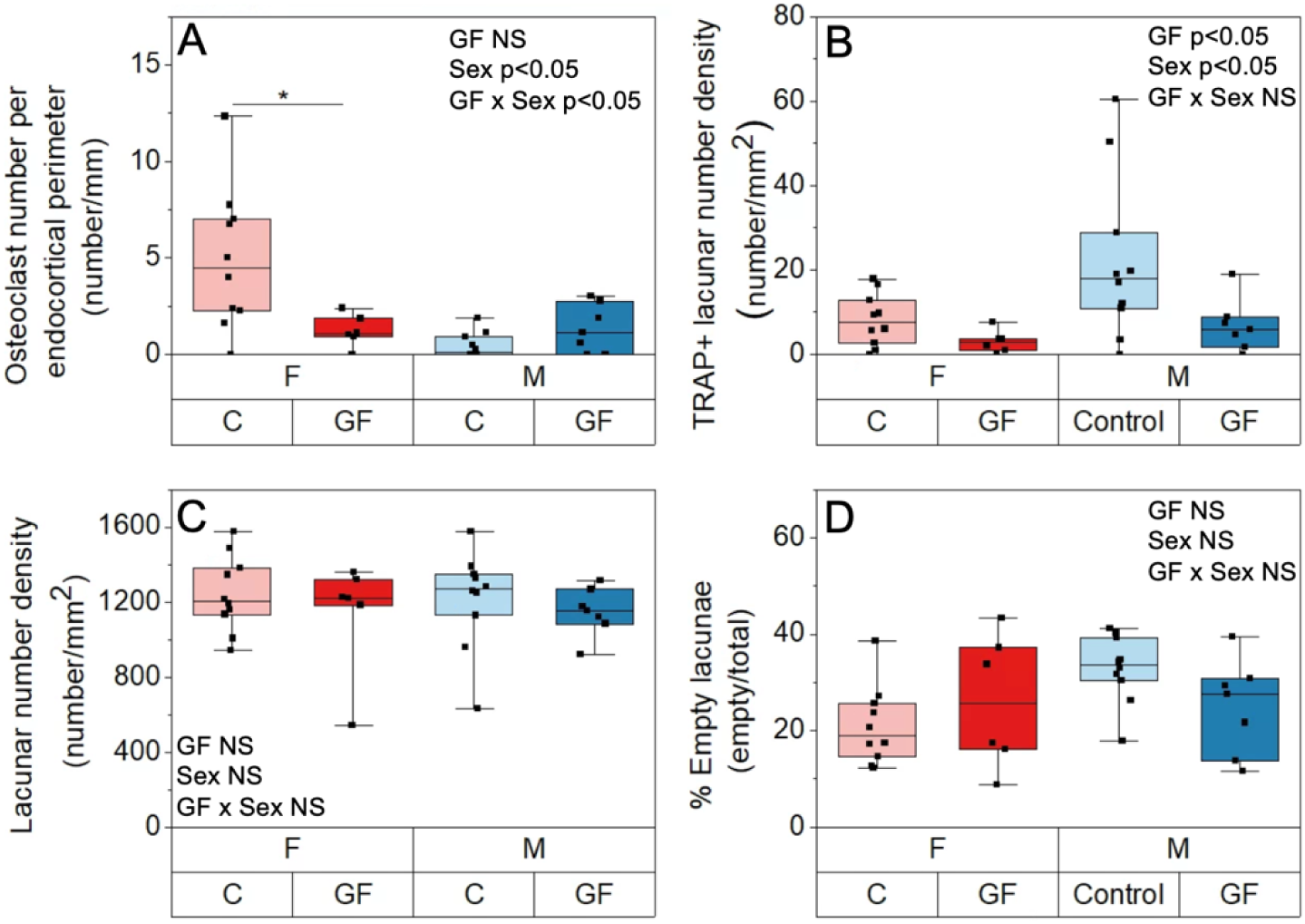
The effect of GF and sex on bone resorbing cells. A) Osteoclast number density per perimeter (number/mm). B) TRAP-stained osteocyte lacunar number density per area (number/mm^2^), C) lacunar number density per area (number/mm^2^), D) percent empty lacunae. Boxplots represent median value (cross), interquartile range (box), minimum/maximum (whiskers), and symbols representing all data points. Significant interactions between sex and GF are shown with asterisks.

### 3.4 Microbiome and sex independently affect bone formation but interactively affect bone resorption

Serum P1NP was higher in GF mice compared to conventional mice (+19.9%, p = 0.03) and in males compared to females (+20.7%, p = 0.008) (**Figure 5A**). Serum CTX1 had a significant interaction between GF and sex (p = 0.036) such that CTX1 level was similar among GF and conventional males but lower and more homogenous in GF females compared with conventional females (-29.7%, p =0.019) (**Figure 5B**). The CTX1/P1NP ratio was lower with GF in both sexes (-26.3%, p = 0.001) and was also higher in females compared to males (+57.6%, p < 0.001) (**Figure 5C**).

**Figure 5.**
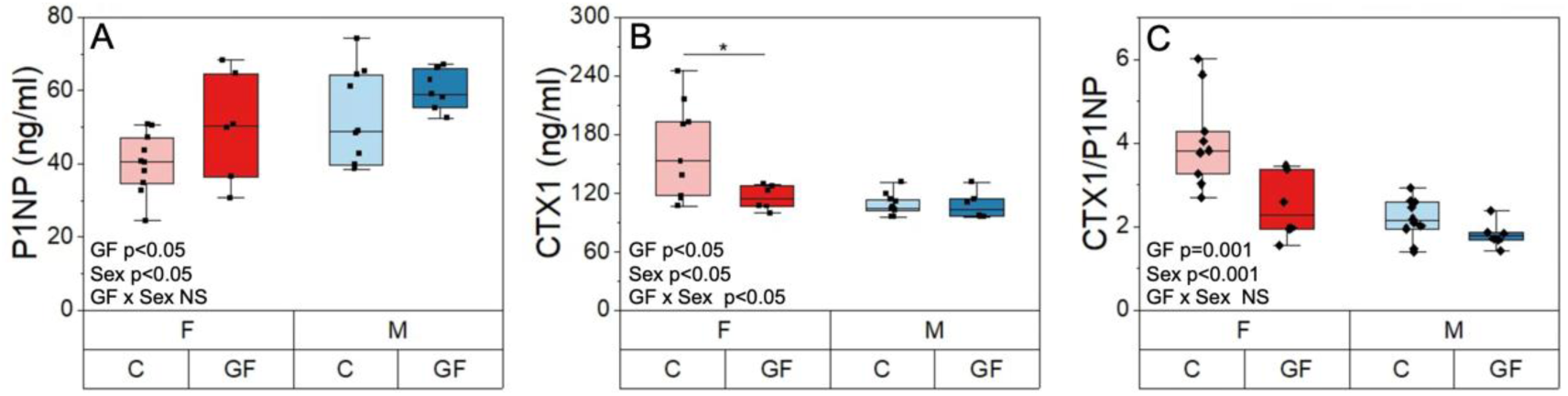
The effect of GF and sex on serum biomarkers of bone turnover. A) P1NP, a biomarker of global bone formation, B) CTX1, a biomarker of global bone resorption, and C) CTX1/P1NP ratio. Boxplots represent median value (cross), interquartile range (box), minimum/maximum (whiskers), and symbols representing all data points. Significant interactions between sex and GF are shown with asterisks.

Both sex and GF influenced alizarin mineralizing surface (MS/BS) at the midshaft femur (**Table 4**). For the periosteal surface, GF mice had higher MS/BS values compared to the conventional mice (+50.6%, p = 0.005). Females had higher MS/BS values compared to males for both periosteal and endocortical surfaces (+29.2%, p = 0.04 and +36.1%, p = 0.046, respectively).

**Table 4.** Histomorphometric analysis of cortical midshaft femurs

| Measure | Female |  | Male |  |
| --- | --- | --- | --- | --- |
|  | Conventional<br>n = 9 | GF<br>n = 5 | Conventional<br>n = 9 | GF<br>n = 4 |
| **Ps.MS/BS (%)**<br>GF: $p = 0.005$<br>Sex: $p = 0.04$<br>GF x sex: NS | 24.0 ± 10.3 | 42.5 ± 7.9 | 22.7 ± 15.5 | 31.3 ± 12.8 |
| **Ec.MS/BS (%)**<br>GF: NS<br>Sex: $p = 0.046$<br>GF x sex: NS | 51.8 ± 19.3 | 55.5 ± 18.9 | 40.9 ± 18.9 | 34.4 ± 15.1 |

Data are presented as mean ± standard deviation.

### 3.5 Microbiome-sex interaction affects cortical geometry and trabecular microstructure

GF increased trabecular bone microstructure, but the effect was more pronounced for males (**Table 5**). BV/TV was higher in GF mice compared to conventional mice (+17.4%, p = 0.05) and was lower in females compared to males (-63.1%, p < 0.001). Similarly, Tb.BMD increased with GF (+9.6%, p = 0.05) and was lower in females compared to males (-41.0%, p < 0.001). Structural modulus index showed more rod-like trabeculae for females (SMI∼3) and more plate-like in males (SMI∼1.5), but GF did not affect this measure. GF and sex had an interactive effect on connectivity density, such that Conn.D greatly increased for GF males (+79.1%, p < 0.001) but remained unchanged for GF females compared to their respective conventional mice. GF and sex had also an interactive effect on Tb.Th, Tb.N and Tb.Sp, such that GF only affected these measures in males and not females. Tb.Th and Tb.Sp were lower in GF males compared to their conventional mice (-13.0%, p = 0.001 and -22.9%, p < 0.001, respectively). Tb.N was higher in GF males compared to their conventional mice (+25.6%, p < 0.001).

**Table 5.**
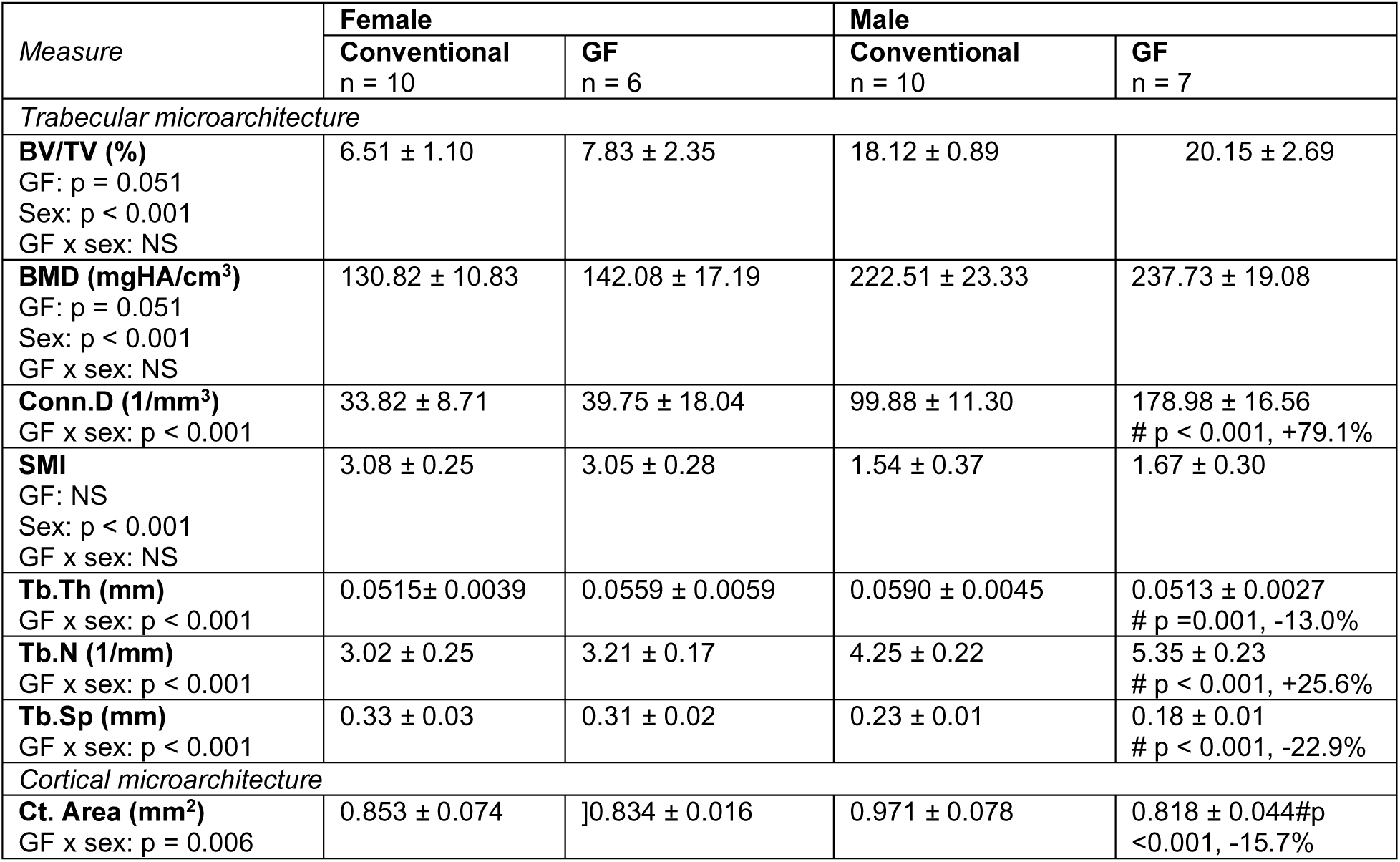
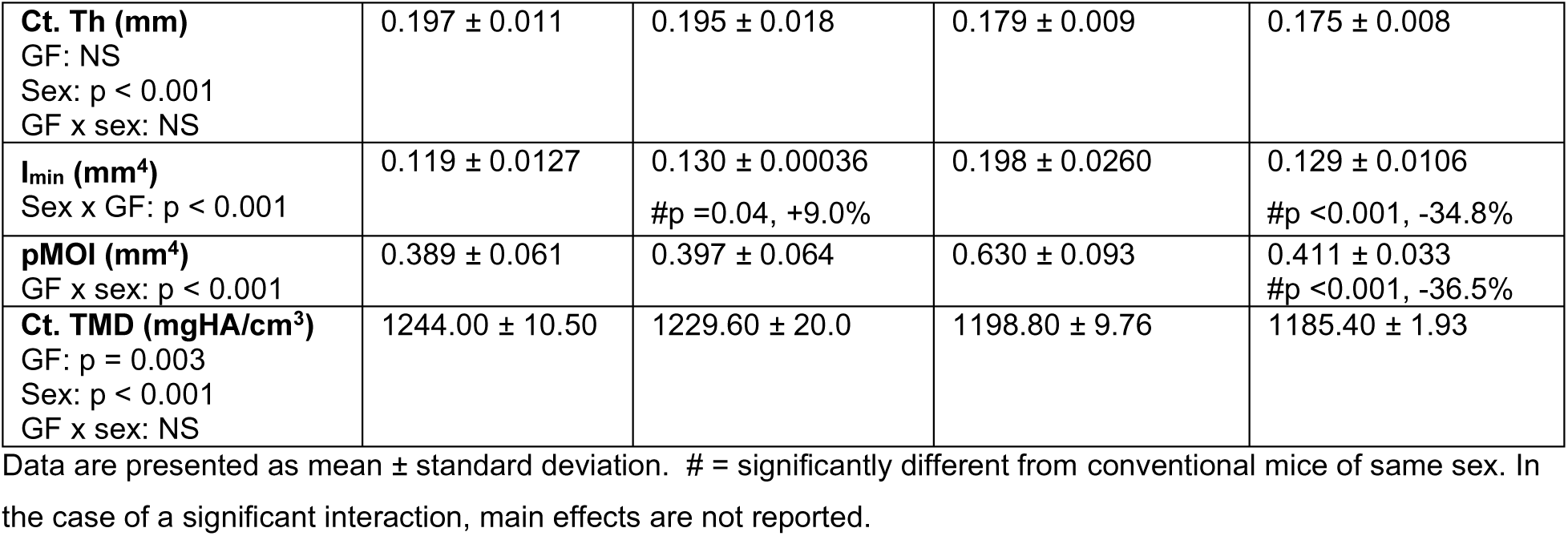
Trabecular microstructure and cortical geometry from microCT analysis

| Measure | Female |  | Male |  |
| --- | --- | --- | --- | --- |
|  | Conventional<br>n = 10 | GF<br>n = 6 | Conventional<br>n = 10 | GF<br>n = 7 |
| *Trabecular microarchitecture* |  |  |  |  |
| **BV/TV (%)**<br>GF: $p = 0.051$<br>Sex: $p < 0.001$<br>GF x sex: NS | 6.51 ± 1.10 | 7.83 ± 2.35 | 18.12 ± 0.89 | 20.15 ± 2.69 |
| **BMD (mgHA/cm<sup>3</sup>)**<br>GF: $p = 0.051$<br>Sex: $p < 0.001$<br>GF x sex: NS | 130.82 ± 10.83 | 142.08 ± 17.19 | 222.51 ± 23.33 | 237.73 ± 19.08 |
| **Conn.D (1/mm<sup>3</sup>)**<br>GF x sex: $p < 0.001$ | 33.82 ± 8.71 | 39.75 ± 18.04 | 99.88 ± 11.30 | 178.98 ± 16.56<br># $p < 0.001$, +79.1% |
| **SMI**<br>GF: NS<br>Sex: $p < 0.001$<br>GF x sex: NS | 3.08 ± 0.25 | 3.05 ± 0.28 | 1.54 ± 0.37 | 1.67 ± 0.30 |
| **Tb.Th (mm)**<br>GF x sex: $p < 0.001$ | 0.0515 ± 0.0039 | 0.0559 ± 0.0059 | 0.0590 ± 0.0045 | 0.0513 ± 0.0027<br># $p = 0.001$, -13.0% |
| **Tb.N (1/mm)**<br>GF x sex: $p < 0.001$ | 3.02 ± 0.25 | 3.21 ± 0.17 | 4.25 ± 0.22 | 5.35 ± 0.23<br># $p < 0.001$, +25.6% |
| **Tb.Sp (mm)**<br>GF x sex: $p < 0.001$ | 0.33 ± 0.03 | 0.31 ± 0.02 | 0.23 ± 0.01 | 0.18 ± 0.01<br># $p < 0.001$, -22.9% |
| *Cortical microarchitecture* |  |  |  |  |
| **Ct. Area (mm<sup>2</sup>)**<br>GF x sex: $p = 0.006$ | 0.853 ± 0.074 | 0.834 ± 0.016 | 0.971 ± 0.078 | 0.818 ± 0.044#<br>$p < 0.001$, -15.7% |
| **Ct. Th (mm)**<br>GF: NS<br>Sex: $p < 0.001$<br>GF x sex: NS | $0.197 \pm 0.011$ | $0.195 \pm 0.018$ | $0.179 \pm 0.009$ | $0.175 \pm 0.008$ |
| **I<sub>min</sub> (mm<sup>4</sup>)**<br>Sex x GF: $p < 0.001$ | $0.119 \pm 0.0127$ | $0.130 \pm 0.00036$<br>#$p = 0.04$, +9.0% | $0.198 \pm 0.0260$ | $0.129 \pm 0.0106$<br>#$p < 0.001$, -34.8% |
| **pMOI (mm<sup>4</sup>)**<br>GF x sex: $p < 0.001$ | $0.389 \pm 0.061$ | $0.397 \pm 0.064$ | $0.630 \pm 0.093$ | $0.411 \pm 0.033$<br>#$p < 0.001$, -36.5% |
| **Ct. TMD (mgHA/cm<sup>3</sup>)**<br>GF: $p = 0.003$<br>Sex: $p < 0.001$<br>GF x sex: NS | $1244.00 \pm 10.50$ | $1229.60 \pm 20.0$ | $1198.80 \pm 9.76$ | $1185.40 \pm 1.93$ |

Data are presented as mean ± standard deviation. # = significantly different from conventional mice of same sex. In the case of a significant interaction, main effects are not reported.

The effect of GF on bone cortical geometry was different between males and females. Cortical area and I_min_ showed significant interactions between sex and GF (p = 0.006 and p < 0.001, respectively) such that male bones were smaller with GF (Ct.Ar: -15.9%, p < 0.001; I_min_: - 34.8%, p < 0.001). GF females had similar Ct.Ar but increased I_min_ (+15.0%, p = 0.04) compared to their conventional mice. Cortical thickness increased with female sex (+10.7%, p < 0.001). Sex and GF also had an interactive effect on polar moment of inertia. pMOI was lower in GF males compared to their conventional mice (+34.7%, p < 0.001) but unchanged among females. GF and sex each affected cortical tissue mineral density. Ct.TMD was slightly lower with GF (- 1.2%, p = 0.003) and female sex (+3.8%, p < 0.001).

### 3.6 Microbiome and sex independently and interactively affect bone strength and modulus but not toughness

The maximum strength from three-point bending was greater for females than males (+15.5%, p < 0.001) and for GF mice than conventional mice (+13.0%, p < 0.001) (**Figure 6A**). The yield strength was greater for females than males (+18.5%, p = 0.005) but not affected by GF or it’s interaction with sex. Modulus showed an interactive effect of GF and sex (p = 0.034) (**Figure 6B**). Modulus was greater for GF males compared to conventional males (+22.7%, p = 0.02), but was not different for GF and conventional females. Toughness from three-point bending test (**Figure 6C**) and K_c-max_ (**Figure 6D**) and K_c-yield_ (**Supp. Table 1**) values from notched fracture test did not differ with GF or sex.

**Figure 6.**
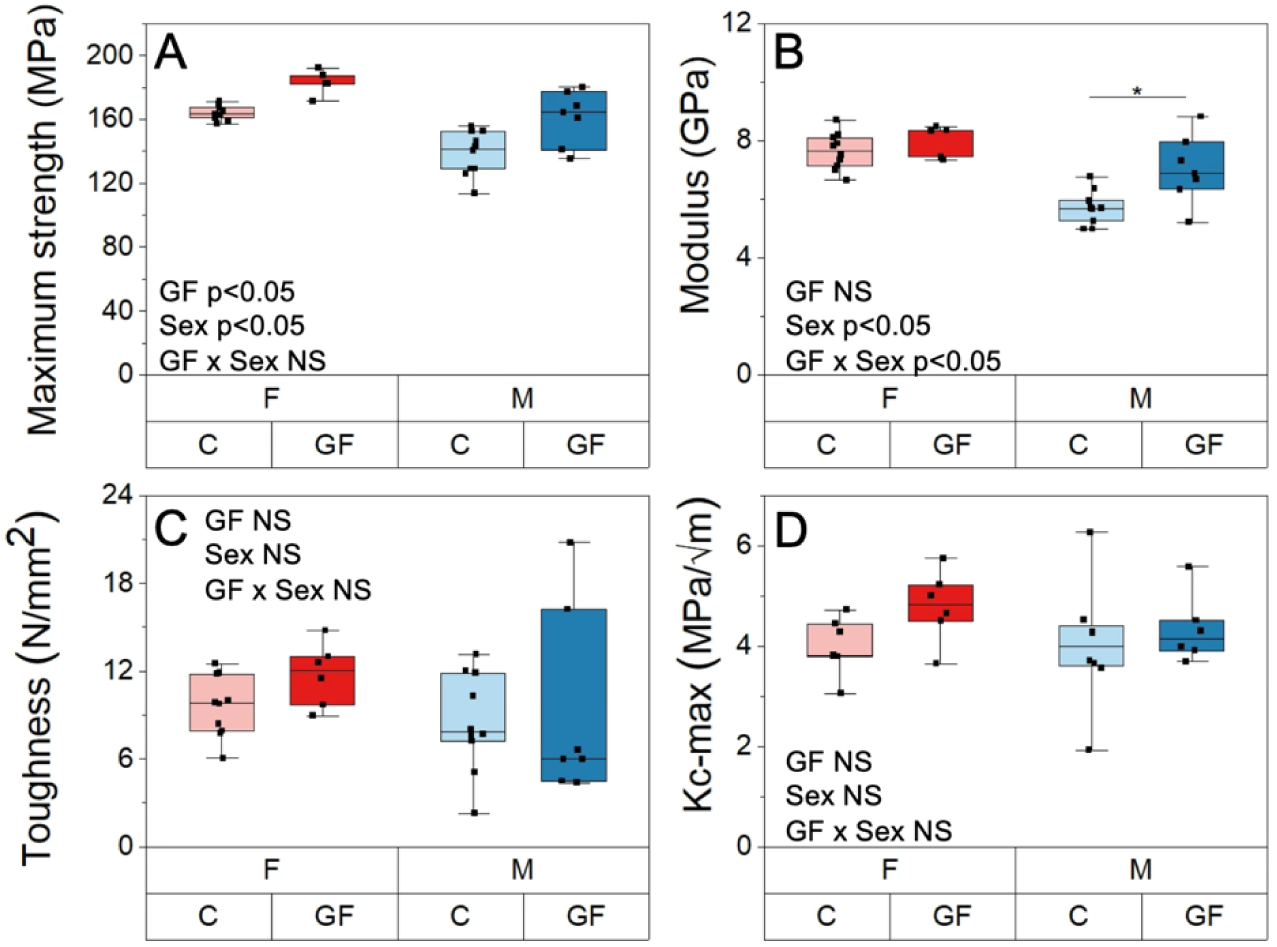
The effect of GF and sex on whole bone material properties. A) Maximum strength from three-point bending, B) modulus from three-point bending, C) toughness from three-point bending, and D) maximum-load fracture toughness (K_c-max_) from notched three-point bending. Boxplots represent median value (cross), interquartile range (box), minimum/maximum (whiskers), and symbols representing all data points. Significant interactions between sex and GF are shown with asterisks.

### 3.7 Microbiome independently affect tissue scale bone mineralization and microbiome-sex interaction affects tissue modulus and cortical porosity

Ca_Peak_ values from qBSE were slightly higher (+3.0%, p = 0.018) with GF but were not affected by sex (**Figure 7A**). Ca_Width_ values were similar among all groups. GF increased the mean nanoindentation modulus only for males (+8.4%, p = 0.023, **Figure 7B**). The standard deviation of *E_i_* was unaffected by GF or sex. Cortical bone porosity estimated from SEM showed an interactive effect of GF and sex (p= 0.023) such that total porosity was decreased for GF males compared to conventional males (-19.6%, p =0.019) but was unchanged for GF females versus conventional females (**Figure 8A**). Lacunar porosity and vascular porosity were not different among groups (**Figure 8C-D**). Pore number density was also not different among groups (**Figure 8D**).

**Figure 7.**
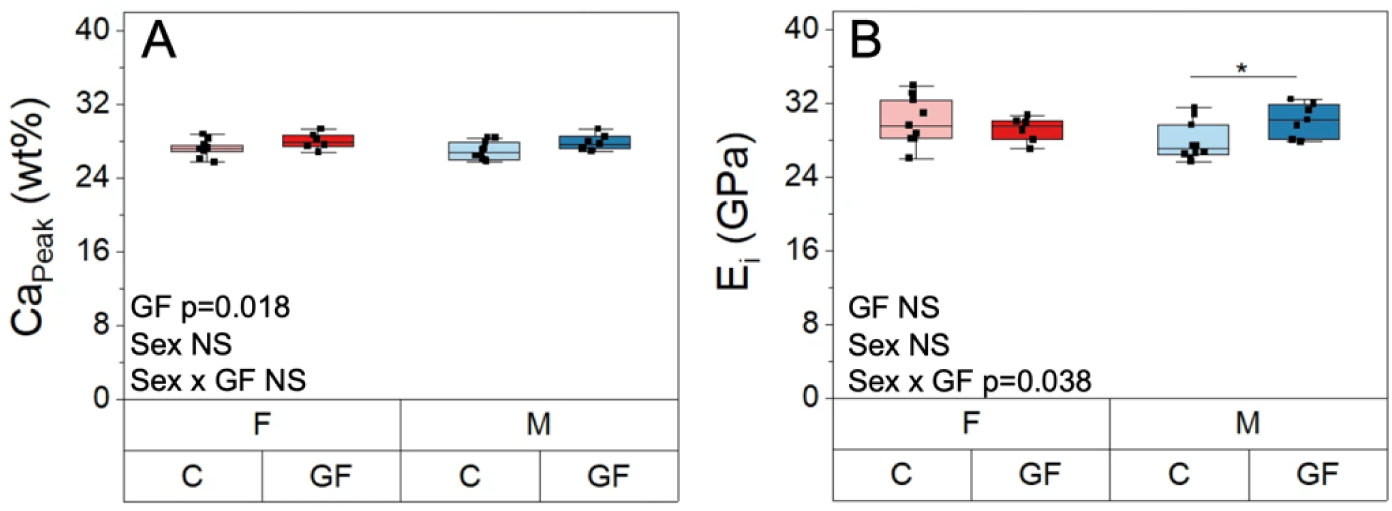
The effect of GF and sex on tissue scale material properties. A) Ca_Peak_ from qBSE, and B) nanoindentation modulus (E_i_). Boxplots represent median value (cross), interquartile range (box), minimum/maximum (whiskers), and symbols representing all data points. Significant interactions between sex and GF are shown with asterisks.

**Figure 8.**
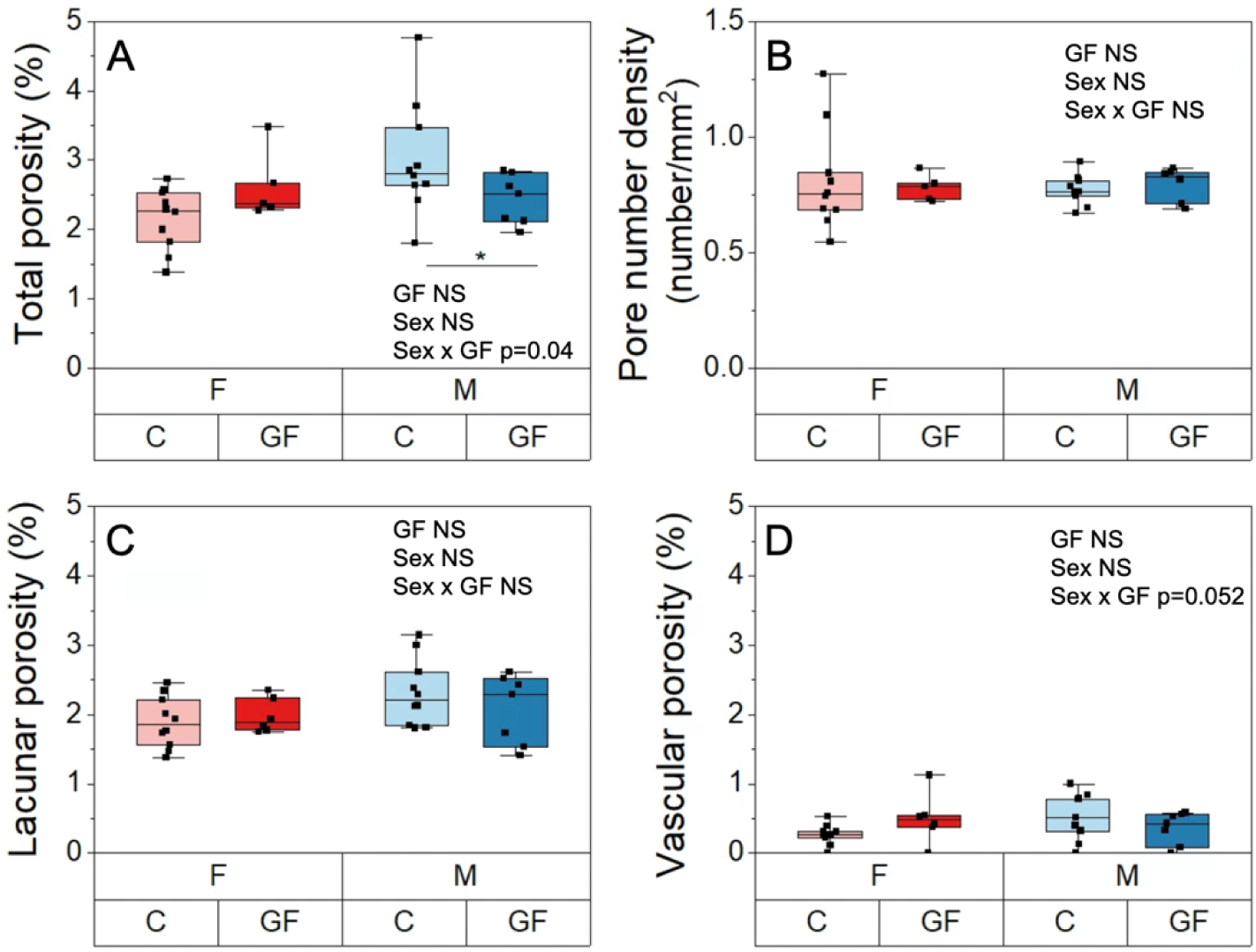
Cortical porosity assessments. A) Total porosity, B) lacunar porosity, C) vascular porosity, and D) pore number density. Boxplots represent median value (cross), interquartile range (box), minimum/maximum (whiskers), and symbols representing all data points. Significant interactions between sex and GF are shown with asterisks.

### 3.8 Microbiome affects mineral-to-matrix ratio and collagen structure but does not affect mineral maturity

Mineral-to-matrix ratio (ν_2_PO_4_ / Amide ΙΙΙ) was higher in GF mice compared to conventional mice for both sexes (+6%, p = 0.024) (**Figure 9A**). Mineral maturity, as indicated by carbonate substitution (ν_1_CO_3_ / ν_1_PO_4_) and crystallinity, was not affected by GF or sex (**Figure 9B**). GF affected Amide I subpeaks intensity, which represents the collagen structure and helical status. Disruption in helical status of the collagen indicates a transition from an ordered triple helical structure to less-ordered forms of structures in collagen^(59)^. The amide I sub-band ratio I_1670_/I_1690_ was higher with GF (+6.6%, p = 0.026). The amide I sub-band ratio I_1670_/I_1640_ was slightly higher with GF (+3.5%, p = 0.053) while I_1670_/I_1610_ was more increased with GF (+8.3%, p = 0.011) (**Figure 9C-D**). Sex did not impact any measures of amide I subpeaks. There were no interactions between sex and GF on Raman measurements of bone composition.

**Figure 9.**
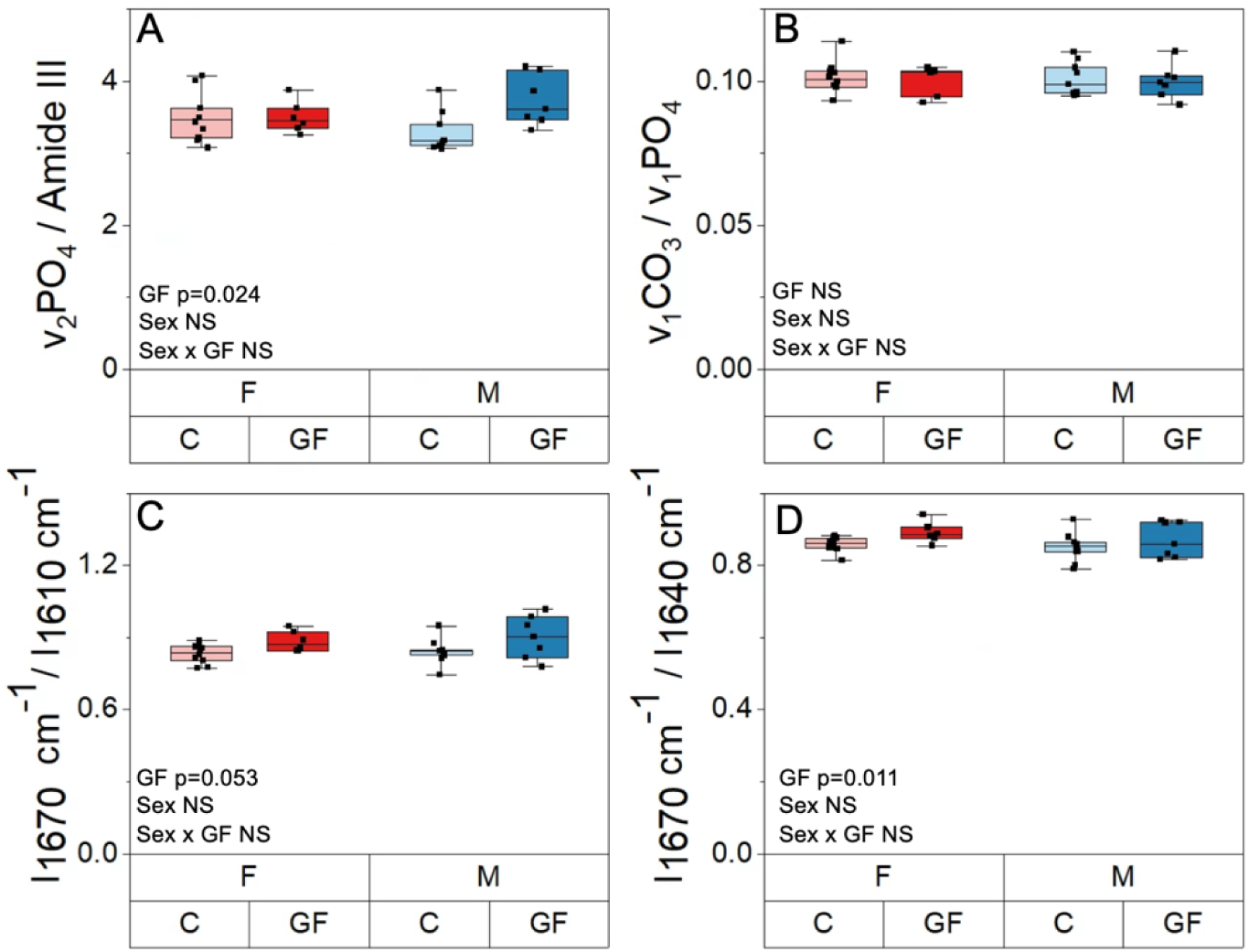
The effect of GF and sex on tissue scale composition. A) Mineral-to-matrix ratio, B) carbonate substitution, C) I_1670_/I_1640_ ratio, and D) I_1670_/I_1610_ ratio. Boxplots represent median value (cross), interquartile range (box), minimum/maximum (whiskers), and symbols representing all data points.

3.9 Sex but not microbiome affects marrow adiposity

Bone marrow adiposity analysis revealed differences in adiposity between mice that differed by sex and GF (**Supp. Table 1**). Mean marrow cavity area was lower with GF compared to conventional mice (-21%, p = 0.001) and for females compared to males (-20%, p = 0.001). Adipocyte counts (-25%, p = 0.02) were decreased with GF and higher and more heterogenous in females compared to males (+359%, p < 0.001). Consequently, adipocyte number density (number of adipocytes per marrow area) was similar between GF and conventional mice and higher and more heterogenous in females compared to males (+452%, p < 0.001). Adipocyte size did not differ with GF nor sex.

3.10 Microbiome and sex each distinctly influence the cortical bone metabolome

A total of 2,129 metabolite features were detected across all humerus cortical bone samples (**Supplement 2**). We used PLS-DA to compare the effects of GF (GF vs. conventional), sex within treatment (GF males vs. GF females, conventional males vs. conventional females), and the effect of GF within sex (GF females vs. conventional females, GF males vs. conventional males). PLS-DA analysis of all four groups displayed minimal overlap, suggesting the metabolomes of all four groups were distinct (**Figure 10A**). Distinct separations were observed between male and female metabolites for GF mice (**Figure 10B**), GF males vs. conventional males (**Figure 10C**), and GF females vs conventional females (**Figure 10D**).

**Figure 10.**
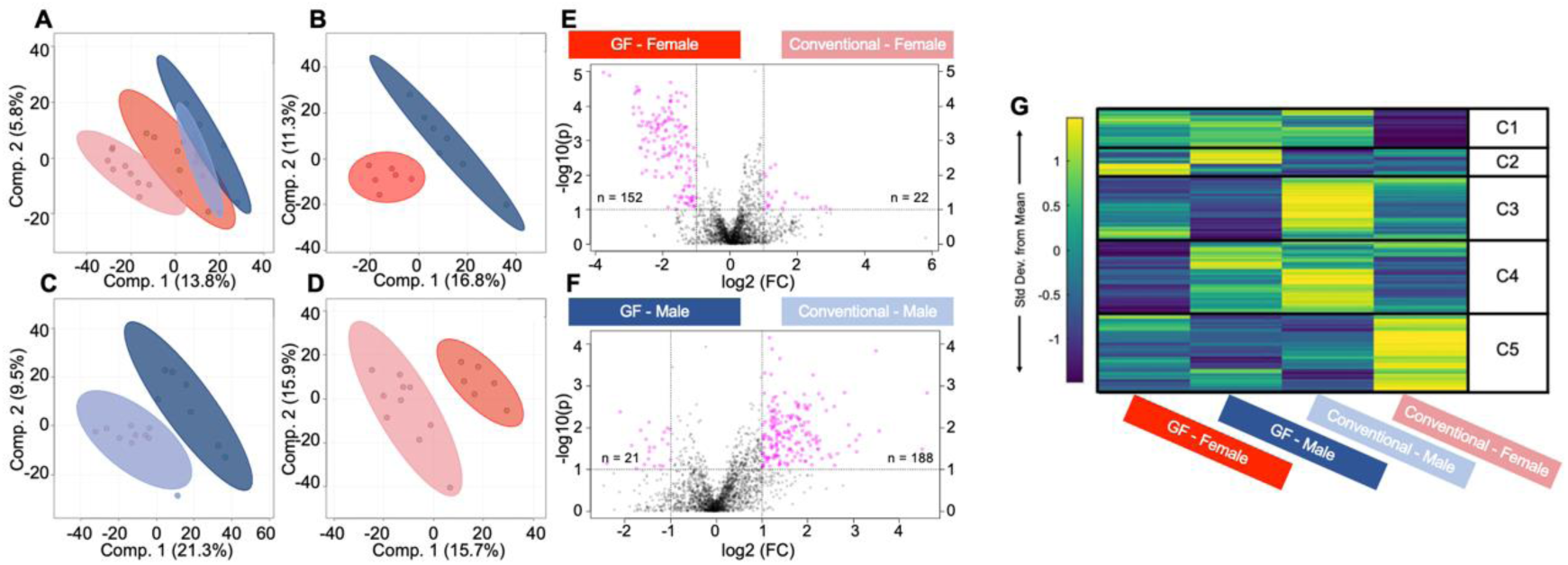
Global metabolomic profiles of humeri derived cortical bone vary by sex and microbiome, as identified by multiple analyses. From supervised PLS-DA analysis, A) metabolites of all four groups showed a clear separation. B) GF males and GF females, C) GF males and conventional males, and D) GF females and conventional females all had distinct metabolomes. From volcano plot analysis (E) metabolite features detected for GF females were significantly different from the ones detected for conventional females, and similarly (F) metabolite features were differently regulated between GF males and conventional males. (G) Median intensity heatmap analysis displayed clusters of metabolites that are differentially regulated between all groups. C1-C5 = clusters 1 – 5.

Volcano plot analysis was utilized to identify sub-populations of metabolite features that were different between GF and conventional for mice of the same sex. In total, 152 features were statistically significant and had higher concentrations in GF females compared to conventional females, whereas 22 metabolite features were statistically significant and had higher concentrations in conventional females compared to GF females. (**Figure 10E**). 188 metabolite features were statistically significant and had higher concentrations in conventional males compared to GF males, whereas 21 were statistically significant and had higher concentrations in GF males compared to conventional males (**Figure 10E**).

Heatmap analysis identified clusters of metabolite features specific to the four groups of male and female GF and conventional mice (**Figure 10G, Supp. Table 2**). Pathways associated with the selected clusters from heatmaps and with the metabolite features from volcano plot were identified for each group. A shared metabolic theme among all females, GF and conventional, was increased levels of glycosaminoglycan degradation. The most significant metabolite features for GF females were increased lipid metabolism (sphingolipid metabolism and arachidonic acid metabolism), whereas for conventional females, significant metabolite features corresponded to increased levels of glycosylphosphatidylinositol (GPI)-anchor biosynthesis and amino acid metabolism (cysteine, methionine). The shared metabolic theme among all males, GF and conventional, was increased levels of amino acid metabolism (alanine, aspartate, glutamate, arginine, histidine, cysteine, methionine). Metabolite features increased in GF males corresponded to increased levels of porphyrin metabolism, whereas features increased in conventional males corresponded to increased levels of purine metabolism, terpenoid backbone biosynthesis, and the pentose phosphate pathway.

### 3.11 Alterations in whole bone quality with microbiome status are multifactorial

The correlations between whole bone material properties and tissue mineralization, collagen structure, cortical porosity, and bone turnover were tested using Spearman’s correlation. We found that tissue mineralization from qBSE was correlated with both strength (Spearman’s ρ = 0.49, p < 0.05) and modulus (ρ = 0.47, p < 0.05). The measure I_1670_/I_1610_ from Raman spectroscopy was also moderately correlated with modulus (ρ = 0.35, p < 0.05) and strength (ρ = 0.32, p = 0.07). Cortical porosity from SEM was correlated with both strength (ρ = -0.38, p < 0.05) and modulus (ρ = -0.45, p < 0.05), but these correlations were evident for males (ρ = -0.57 for strength, ρ = -0.56 for modulus, p < 0.05) and not so much for females (ρ = 0.30 for strength, ρ = 0.20 for modulus, p > 0.05). These independent variables were not correlated or weakly correlated with each other (ρ < 0.3, p > 0.05). Bone strength was not strongly correlated with changes to global bone turnover (CTX1:P1NP, ρ = 0.15, p > 0.05) or osteoclast number density (ρ = 0.25, p > 0.05). Modulus was fairly correlated with CTX1:P1NP and not correlated with osteoclast number density (ρ = 0.35 and p = 0.06, respectively and p > 0.05). Both strength and modulus were correlated with local bone formation in periosteal surface (Ps.MS/BS, ρ = -0.68 for strength, ρ = -0.44 for modulus, p < 0.05) but not with endocortical surface bone formation (ρ < -0.01).

## 4. Discussion

The germ-free (GF) mouse model is expected to increase bone mass (**Table 1**). However, the influence of the gut microbiome on bone quality is not well understood because the effects of GF on bone tissue metabolism, bone cell remodeling, bone tissue mineralization and resulting tissue-scale through whole bone material properties are not known. Also unknown is how the effects of GF on bone differ between sexes. Here, we determined that GF indeed affects bone tissue metabolism, many features of bone cells abundance and remodeling, and multiscale bone quality, and that these effects are for the most part sexually dimorphic.

Our data demonstrate that the effect of the gut microbiome on bone formation and resorption is different for females and males (**Table 6**). For both sexes, GF increased cortical bone formation at the femur diaphysis; however, the increase in bone formation was two-fold greater for GF females compared to GF males. Further, GF females had reduced osteoclast density in cortical bone, but GF males did not. These cortical femur-specific data aligned with global serum biomarkers of bone formation and resorption. Our data demonstrate that the adipocytes density does not change with GF, potentially indicating mesenchymal stem cells differentiation activity may not be influenced by GF status. GF mice have an immature immune system^(18, 24)^, which would be expected to influence precursors available for differentiation to osteoclasts. The presence of sexual dimorphism in immunological responses to different diseases has been previously reported, with greater pro-inflammatory cytokine responses and T-cell proliferation in female human and mice compared to their male counterparts^(62, 63)^.

**Table 6.** Key findings for the influence of GF and sex on bone multiscale properties.

| Measure |  | No interaction between GF and sex |  | Significant interaction between GF and sex |  |
| --- | --- | --- | --- | --- | --- |
|  |  | GF vs. conventional | Female vs. male | GF females vs. conventional females | GF males vs. conventional males |
| *Bone remodeling* |  |  |  |  |  |
| Midshaft femur | Ps.MS/BS | ↑ 50.6%** | ↑ 29.2%* | NS | NS |
|  | Ec.MS/BS | NS | ↑ 36.1%* | NS | NS |
| Serum biomarkers | P1NP | ↑ 19.9%* | ↓ 20.7%** | NS | NS |
|  | CTX1 | - | - | ↓ 29.7%* | NS |
|  | CTX1/P1NP | ↓ 26.3%** | ↑ 57.6%*** | NS | NS |
| Proximal tibia- | % TRAP+ Lac. | ↓ 67%* | ↓ 155%* | NS | NS |
| Histology | Oc. N. density | - | - | ↓ 75%* | NS |
| Flushed tibia-PCR | MMP2 | ↓ 38.9%* | NS | NS | NS |
|  | MMP14 | NS | ↑ 65.2%* | NS | NS |
|  | OPG | ↓ 50.5%* | NS | NS | NS |
|  | RankL/OPG | ↑ 127.6%* | NS | NS | NS |
|  | CTSK | NS | ↑ 97.1%* | NS | NS |
| *Bone microstructure and geometry* |  |  |  |  |  |
| Cancellous: distal | BV/TV | ↑ 17.4%* | ↓ 63.1%*** | NS | NS |
| Femur metaphysis | Conn.D | - | - | NS | ↑ 79.1%*** |
|  | Tb.Th | - | - | NS | ↓ 13.0%** |
| Cortical: midshaft | Ct.Ar | - | - | NS | ↓ 15.9%*** |
| Femur | I<sub>min</sub> | - | - | ↑ 16.0%* | ↓ 34.8%*** |
|  | Ct.TMD | ↓ 1.2%** | ↑ 3.7%*** | NS | NS |
| *Whole material properties* |  |  |  |  |  |
| Femur | Modulus | - | - | NS | ↑ 22.7%* |
|  | Max strength | ↑ 13.0%** | ↑ 15.5%** | NS | NS |
|  | Yield strength | NS | ↑ 18.5%** | NS | NS |
|  | Toughness (3PB) | NS | NS | NS | NS |
|  | K<sub>c-max</sub> and K<sub>c-yield</sub> | NS | NS | NS | NS |
| *Tissue-scale material properties and cortical porosity* |  |  |  |  |  |
| Midshaft femur | E<sub>i</sub> | - | - | NS | ↑ 8.4%* |
|  | Stdev. E<sub>i</sub> | NS | NS | NS | NS |
|  | Ca<sub>Peak</sub> | ↑ 3.0%* | NS | NS | NS |
|  | Ca<sub>Width</sub> | NS | NS | NS | NS |
|  | Cortical porosity | - | - | NS | ↓ 19.6%* |
| *Tissue composition and collagen structure* |  |  |  |  |  |
| Humerus | Mineral/ matrix | ↑ 6.0%* | NS | NS | NS |
|  | Car / Phos | NS | NS | NS | NS |
|  | Crystallinity | NS | NS | NS | NS |
|  | I<sub>1670</sub> / I<sub>1610</sub> | ↑ 8.3%* | NS | NS | NS |
|  | I<sub>1670</sub> / I<sub>1640</sub> | ↑ 3.5%* | NS | NS | NS |
| *Bone metabolism* |  |  |  |  |  |
| Proximal tibia- | Mar. cavity area | ↓ 21%** | ↓ 20%** | NS | NS |
| Histology | Adp. count | ↓ 25%* | ↑ 359%*** | NS | NS |
| Humerus |  | ↑ lipid metabolism in GF females*. |  |  |  |
| Metabolomics |  | ↑ GPI-anchor biosynthesis in conventional females*. |  |  |  |
|  |  | ↑ porphyrin metabolism in GF males*. |  |  |  |
|  |  | ↑ amino acid metabolism in conventional males*. |  |  |  |

\* indicates p < 0.05, \*\* indicates p < 0.01, and \*\*\* indicates p < 0.001. when an interaction exists, main effects of GF and sex are not reported.

Females also have enhanced innate and adaptive immune responses to inflammation or bacteria-driven diseases^(64, 65)^. Similarly, evidence supports sex differences in osteoclast differentiation and precursor population, although specific results are contradictory^(65–68)^. Some studies reported that *in vitro* osteoclastogenesis occurs faster in osteoclast precursor derived from female mice cells compared to male cells in the absence of pathogens^(66, 68)^ while others reported bacterial-induced osteoclastogenesis is faster in *in vitro* male osteoclast precursor cells compared to females^(67)^. The sex differences in the decline of osteoclast number in our study implies a likely difference in either the immune systems of male and female GF mice or a sexual dimorphism in the resilience of osteoclast differentiation on the immune system.

The osteocyte regulates bone remodeling, and prior work suggests that osteoblast to osteocyte transition may be decreased in GF mice through disruptions in immune system and bacteria-derived vitamin K_2_ biosynthesis^(29, 37)^. Therefore, we investigated the influences of GF and sex on osteocyte abundance, gene expression related to osteocyte control of osteoclast and osteoblast differentiation and lacunar-canalicular system remodeling, and osteocyte perilacunar bone resorption. We found that GF status did not alter lacunar number density or percent empty lacunae for either females or males. GF increased RankL/OPG ratio by downregulating OPG expression in both males and females. The RankL-OPG signaling system regulates osteoclastogenesis in the marrow^(69)^ and the downregulation of OPG promotes osteoclastogenesis and osteoclastic activity^(70)^. Osteocyte perilacunar bone resorption, as estimated from TRAP positive lacunae, was decreased with GF status. However, GF status did not impact most measures of gene expression related to lacunar-canalicular system remodeling. These data suggest efforts by osteocytes in the context of the GF model to decrease bone mass and also participate in lacunar-canalicular bone remodeling but failure to achieve reduced osteoblast activity or increased osteoclast activity.

GF affects multiscale bone quality and these changes depend on sex (**Table 6**). We found that bone volume fraction was similarly increased for GF mice of both sexes while other features of bone microarchitecture (e.g., Tb.N, Conn.D) were only increased for males. These data align with prior reports of increased bone mass for GF, which only included females^(10, 18, 21)^. Cortical porosity was decreased with GF but only for males. While whole bone strength similarly increased for females and males, modulus was only increased for males. Importantly, because fracture toughness was not improved with GF, we do not interpret GF as improving bone quality. At the tissue scale, for both sexes, cortical bone mineralization at the femur diaphysis was slightly increased with GF, as measured by qBSE and confirmed with Raman spectroscopy. GF did not affect bone mineral maturity measured with Raman spectroscopy of the hydrated femur periosteal surface. GF altered collagen structure (i.e, I_1670_/I_1610_) for both sexes, indicative of transitioning from a collagen organization with ordered triple helical structure to less-ordered forms^(59)^. GF slightly increased cortical bone nanoindentation modulus for males but not females, demonstrating that the summative effects of increased mineralization and disrupted collagen structure do not culminate in the same impact to bone stiffness for both sexes. These data demonstrate that the gut microbiome influences bone microarchitecture, composition, and material properties, and that the specific impacts are sexually dimorphic.

We sought to investigate whether the increase in whole bone material properties (strength and modulus) with GF was driven by tissue mineralization, collagen structure, or cortical porosity. Several studies reported that changes in bone strength and modulus positively correlate with tissue mineralization^(71–76)^ and with collagen crosslinks maturity, structural stability, and fibril orientation^(77–84)^. It is also well-established that high cortical porosity negatively correlates with bone’s elastic modulus and strength^(75, 76, 85–87)^. We found that bone strength and modulus were each moderately correlated with both tissue mineralization from qBSE and collagen structure for both sexes and with cortical porosity only for males. These factors were not intercorrelated (ρ < 0.3). Therefore, alterations in whole bone quality resulting from GF are likely the result of multiple contributing factors including at least mineralization, collagen structure, and cortical porosity. Because GF increased bone formation and decreased bone resorption, we also asked whether the changes to whole-bone material properties were the result of disrupted bone turnover. We found that whole bone biomechanics for both sexes are not or weakly correlated with measures of global bone turnover and are moderately correlated with measures of local turnover (just Ps.MS/BS and not Ec.MS/BS). These results demonstrate that the impact of the gut microbiome on the whole bone quality is partially, but not fully, determined by impacts on bone remodeling activity. The lack of microbiome can cause several other important developmental differences in the skeleton compared to conventional mice.

These differences include increased bone mass in growing C57BL/6 mice^(18)^, shorter femurs in 7 weeks old male BALB/c mice with smaller and thinner cortical area and lower bone volume fraction^(46)^, and increased cortical thickness in 10-12 weeks old female C57BL/6 mice^(19, 20)^.

Since GF had a sexually dimorphic effect on bone quality and remodeling bone cells, we studied sex differences in how GF affects bone cell metabolism. We evaluated the metabolism of cortical bone, which is predominantly populated by osteocytes. We found that compared to conventional mice of the same sex, female GF mice had increased lipid metabolism (highest of all groups) and male GF mice had differentially regulated metabolites in energy metabolism (i.e., upregulated porphyrin metabolism in GF males and upregulated purine metabolism in conventional males). There was not increased adipocyte number density in GF mice, demonstrating that the increased levels of lipid metabolism are not a consequence of increased differentiation of mesenchymal stem cells to adipocytes. GF females also had increased levels of arachidonic acid metabolites compared to conventional females which are reported to be inhibitors of osteoclastic function^(88)^. Together, these findings potentially indicate osteoclast population and bone resorption activity in GF females could be impacted from altered dynamics of lipid metabolism. Conversely, conventional females had increased levels of cysteine and its precursor methionine compared to GF females^(89)^. Cysteine is a key component of cathepsin k protease that predominantly expressed in osteoclasts, and it is essential to bone resorption activity^(90)^. Conventional females had the highest osteoclast population and global resorption activity among all groups which drastically decreased with GF. Males had increased levels of energy and amino acid metabolisms compared to females, with GF males having the highest levels of porphyrin metabolism and conventional males having the highest levels of purine metabolism. This finding is consistent with GF males having the highest bone formation (P1NP and BV/TV) in all groups, based on our prior finding^(45)^. We have previously reported that in conventional mice, bone cells from males and females rely on different metabolic pathways to meet their energy demands. While cells from male mice used amino acid metabolism, cells from females predominantly utilized lipid metabolism^(45)^. It appears that in GF mice, these differences between male and female cortical bone metabolome become much more predominant. GF females and males both build more bone compared to conventional mice, but this evidence suggests that they may engage in different energy metabolism to do so. These results demonstrate that GF affects the bioenergetics of bone cells and that this impact is different in males and females.

A strength of our study is evaluating sex differences in multiscale bone quality for skeletally-mature mice. Through this work we have accumulated, to our knowledge, the largest dataset of sex differences in bone quality amongst conventional C57BL/6 mice currently available in the literature. Key differences pertaining to whole bone mechanics and microarchitecture include higher bone strength, lower trabecular microstructure and smaller cortical geometry in females compared to males. Sex differences related to bone remodeling include increased bone mineralizing surface, higher bone turnover, decreased osteocyte perilacunar bone resorption, higher osteoclast number density, increased cathepsin K and MMP14 expression, and higher adipocyte number for females. Meanwhile, tissue-scale mineralization and collagen structure were similar for conventional females and males. We anticipate that these reference data will be broadly useful in interpreting sex differences in bone quality occurring in disease models and other interventions.

Our study has several limitations. First, GF mice have developmental differences with conventionally-raised mice^(28, 91)^, which likely have a confounding role in the effects of lack of microbiome on the skeleton. Second, it was necessary to characterize bone at multiple skeletal locations due to different sample preparation requirements for each technique, though skeletal site differences are likely to impact the reported results. Most importantly, GF models are not directly translatable to humans. Nonetheless, GF models provide unique insights about the origin of the microbiome impacts on bone quality that are not accessible from other models.

We conclude that absence of gut microbiome in C57BL/6 mice not only increases bone mass but also impacts multiscale bone quality including the organization and mineralization of the bone tissue composite. However, these impacts do not result in changes in the whole bone fracture toughness and only increase the bone strength. The microbiome impact on bone quality is sex dependent. We report that that GF alters the activities and abundance of remodeling bone cells in a sexually dimorphic manner. These alterations start at the level of the cellular metabolome as we demonstrate that cortical bone metabolome is distinct between GF groups and conventional groups and is also sexually dimorphic. Our findings advance our fundamental understanding about the role of microbiome and sex on bone quality.

## Acknowledgments

We are thankful for assistance in germ-free mouse handling from Mark McAlpine. LC-MS analyses were performed with help from Dr. Don Smith and Dr. Katie Steward and the Montana State University Proteomics, Metabolomics and Mass Spectrometry Facility. Maria Jerome is thanked for preparing histology samples and Dr. Heidi Smith and Dr. Markus Dieser are appreciated for their assistance with microscopy. Dr. Nathaniel Rieders and Dr. Sara Maccagnano-Zacher provided useful advice in collecting qBSE images. The Center for Advanced Orthopaedic Research at Beth Israel Deaconess performed microCT analyses. Brady Hislop assisted with tissue harvest. Funding for this study was provided by the National Institutes of Health (NIGMS P20 GM103474, NIA R03 AG068680, NSF 1554708 and 2140127, and NIAMS 5R01AR073964)

## Supplement 1

**Supp. Figure 1.**
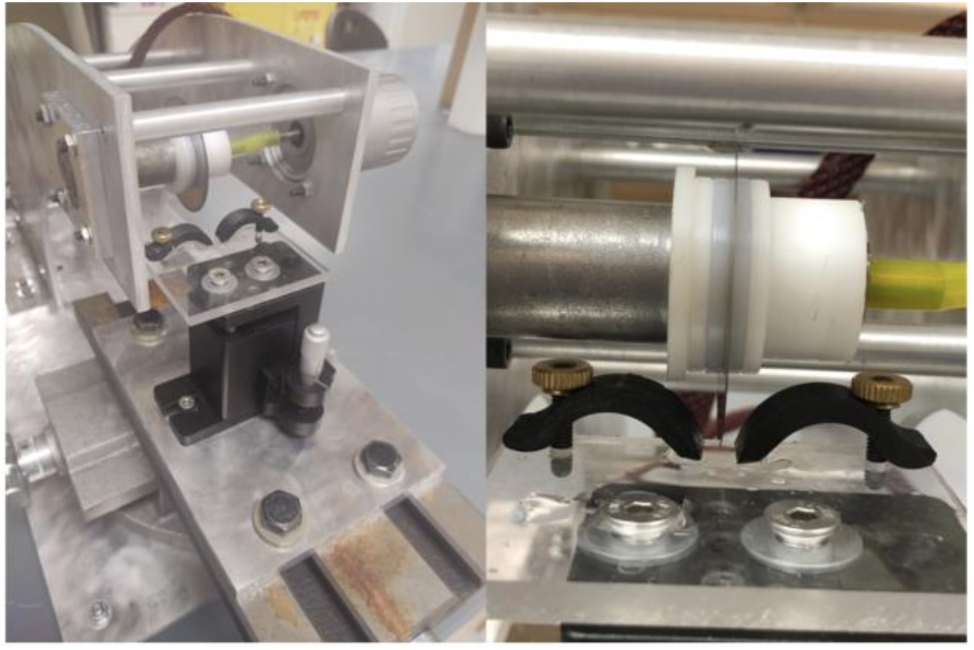
Notch fracture instrument

**Supp. Figure 2.**
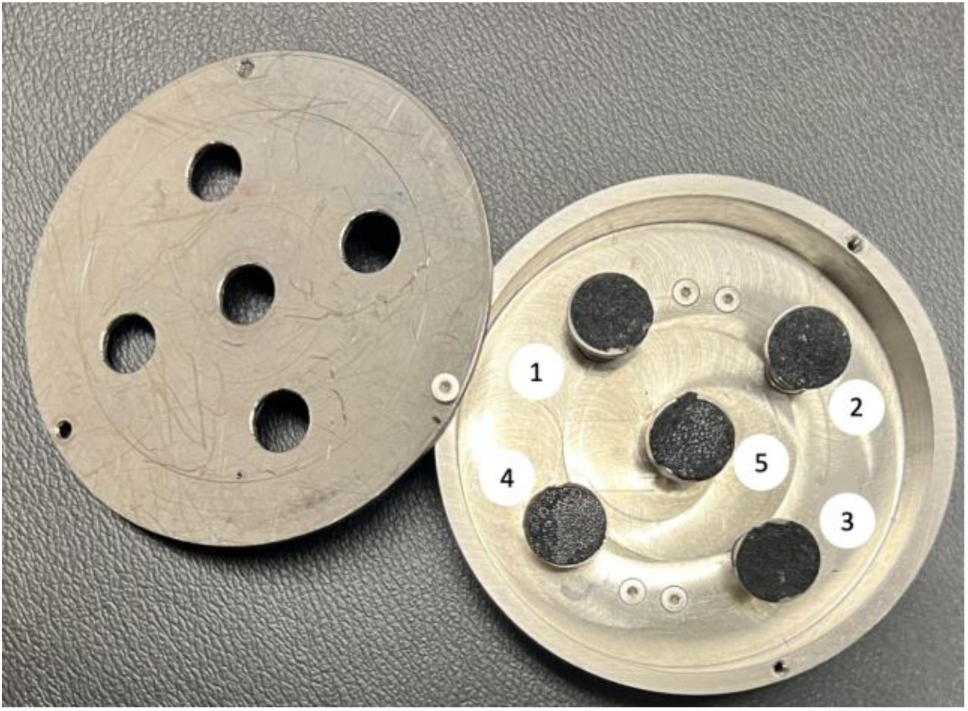
SEM sample holder

**Supp. Table 1.**
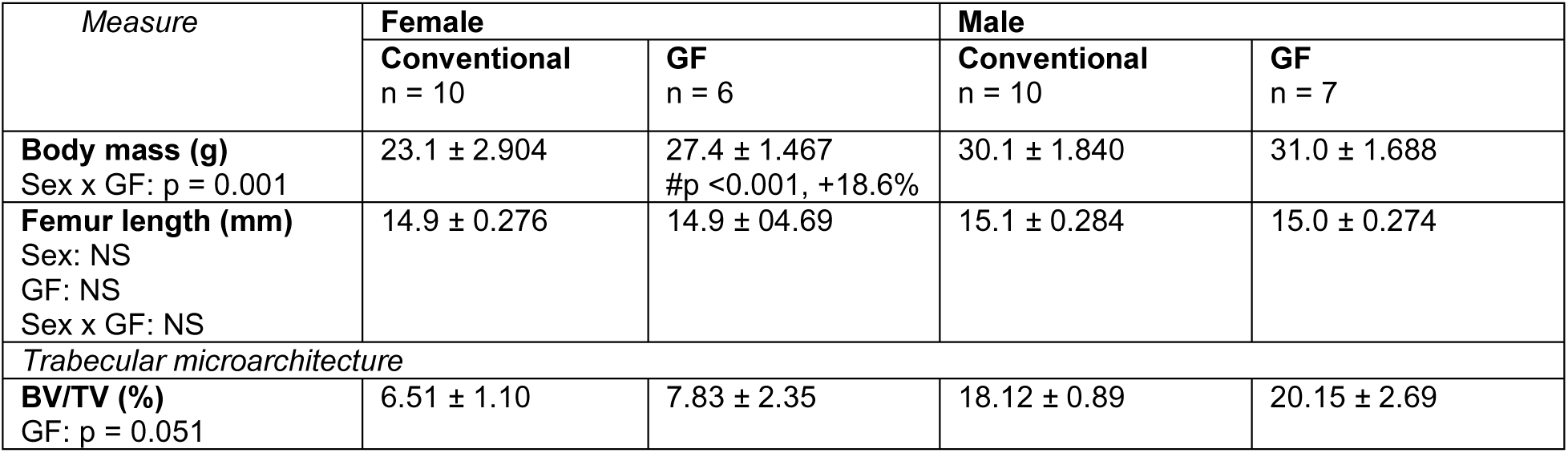
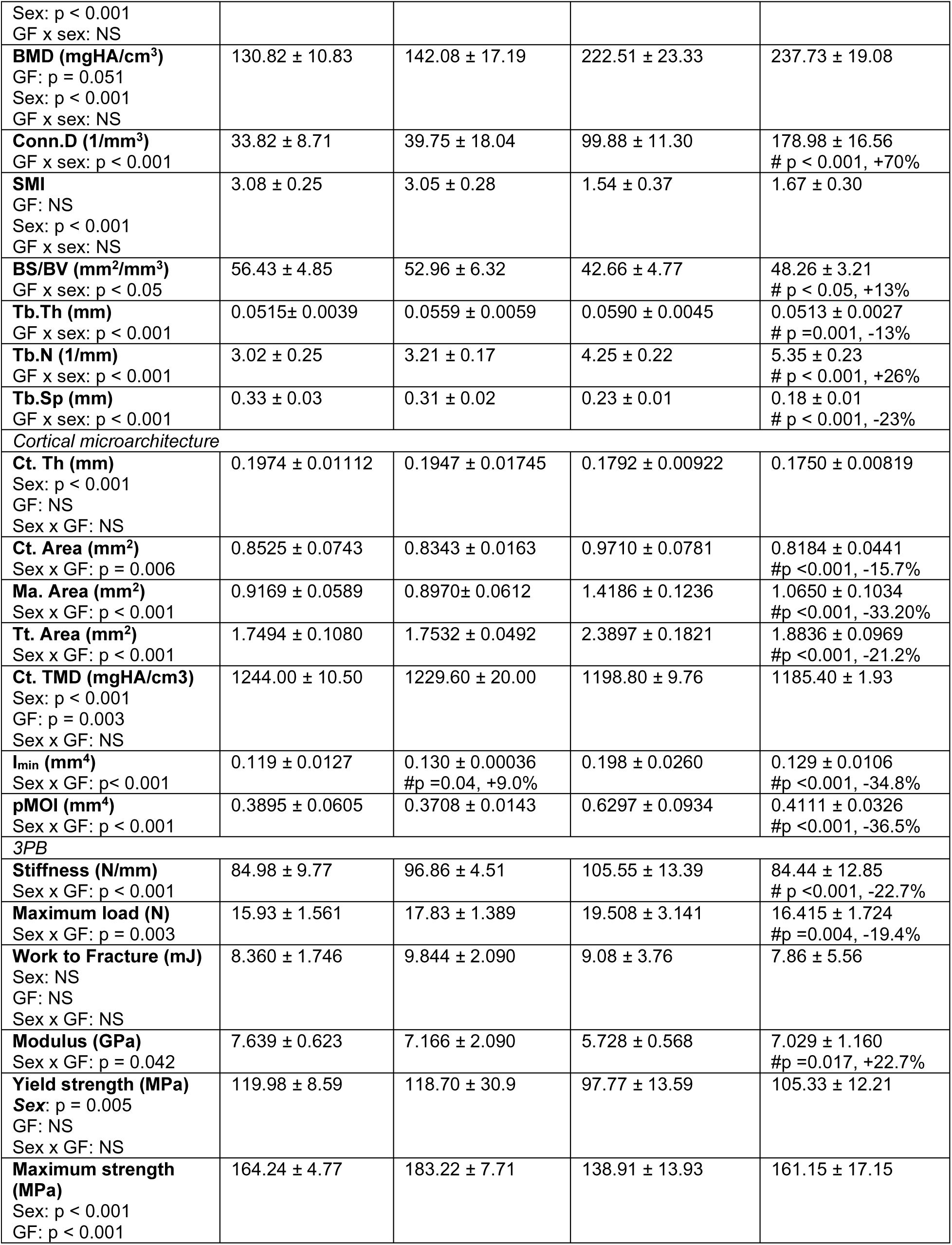
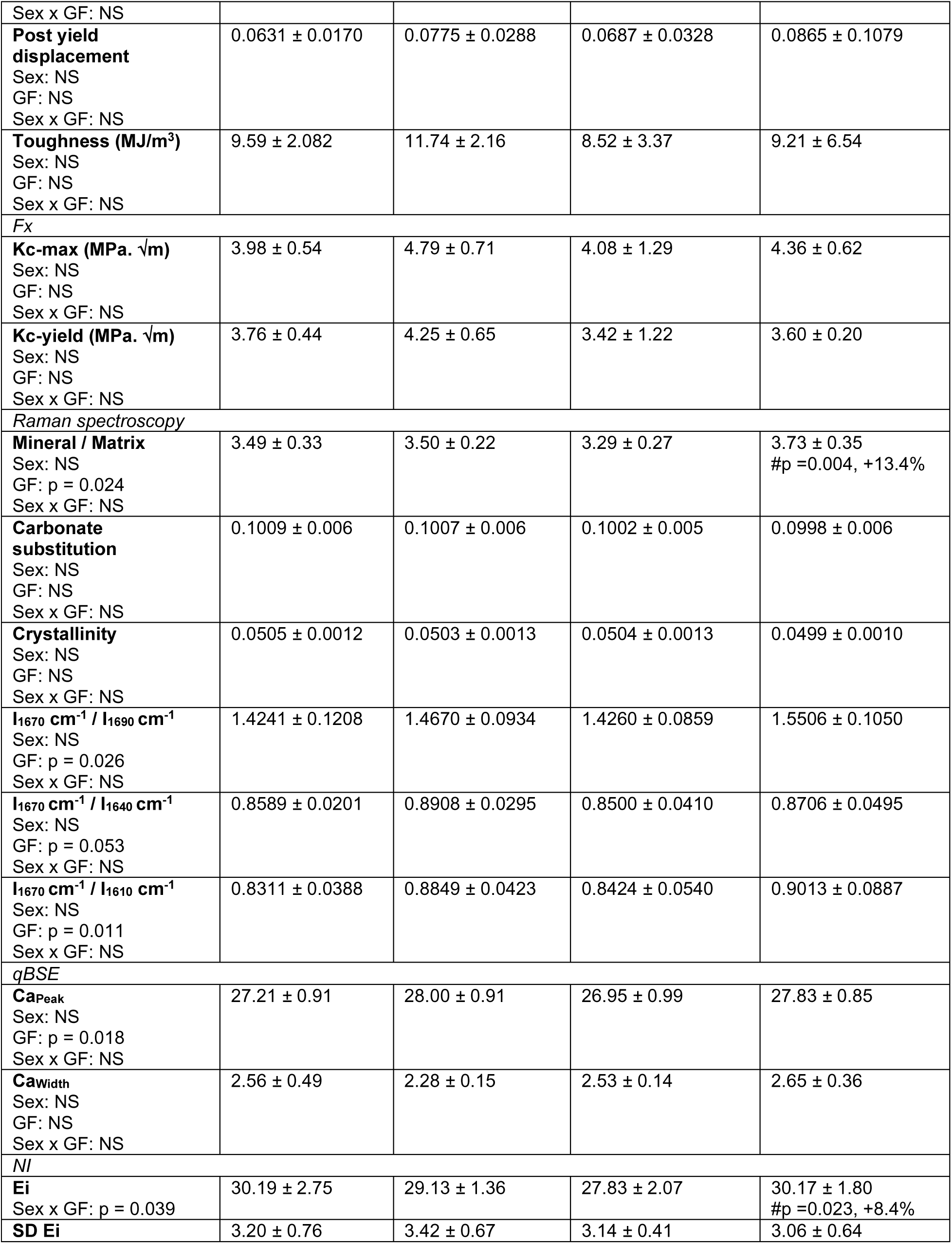
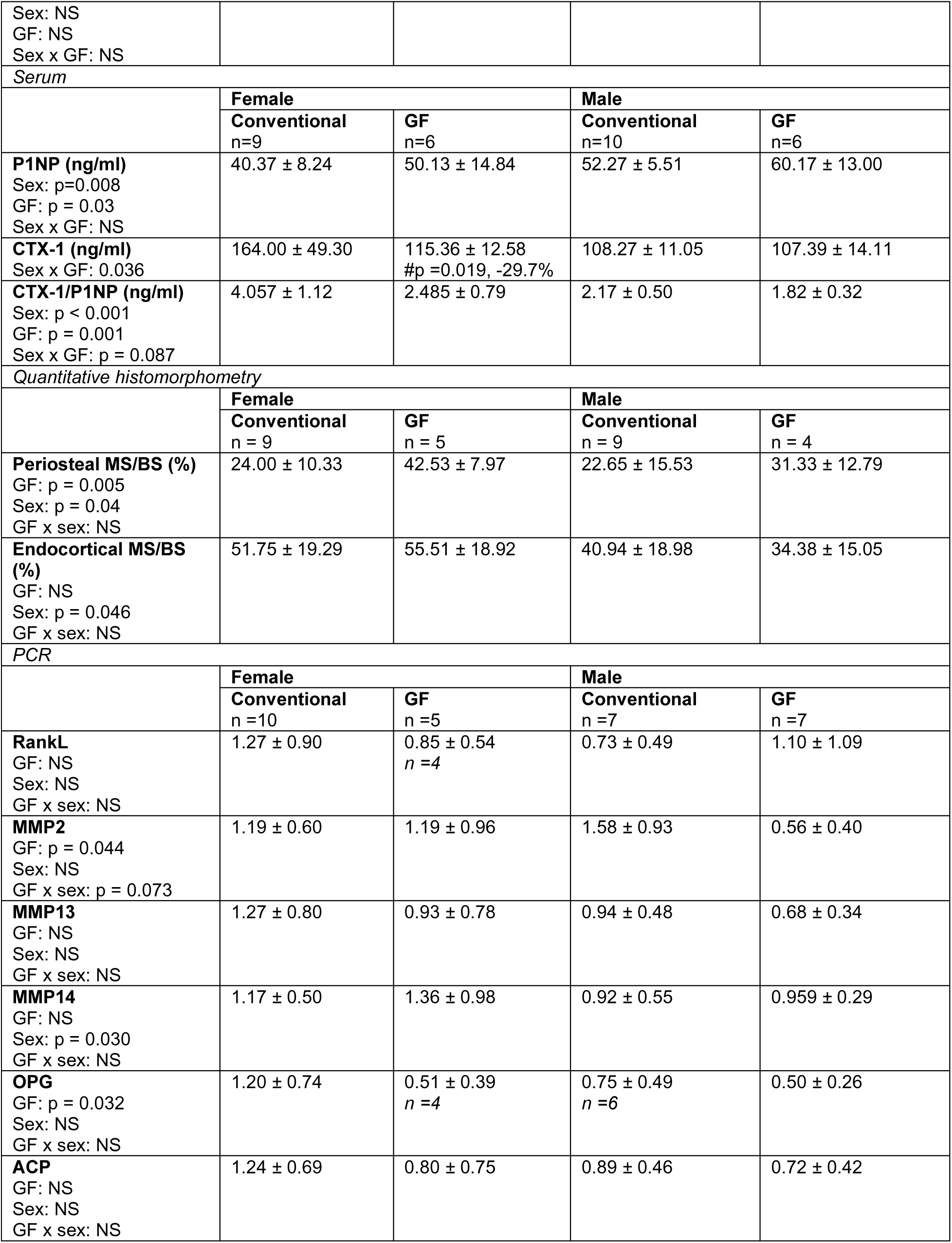
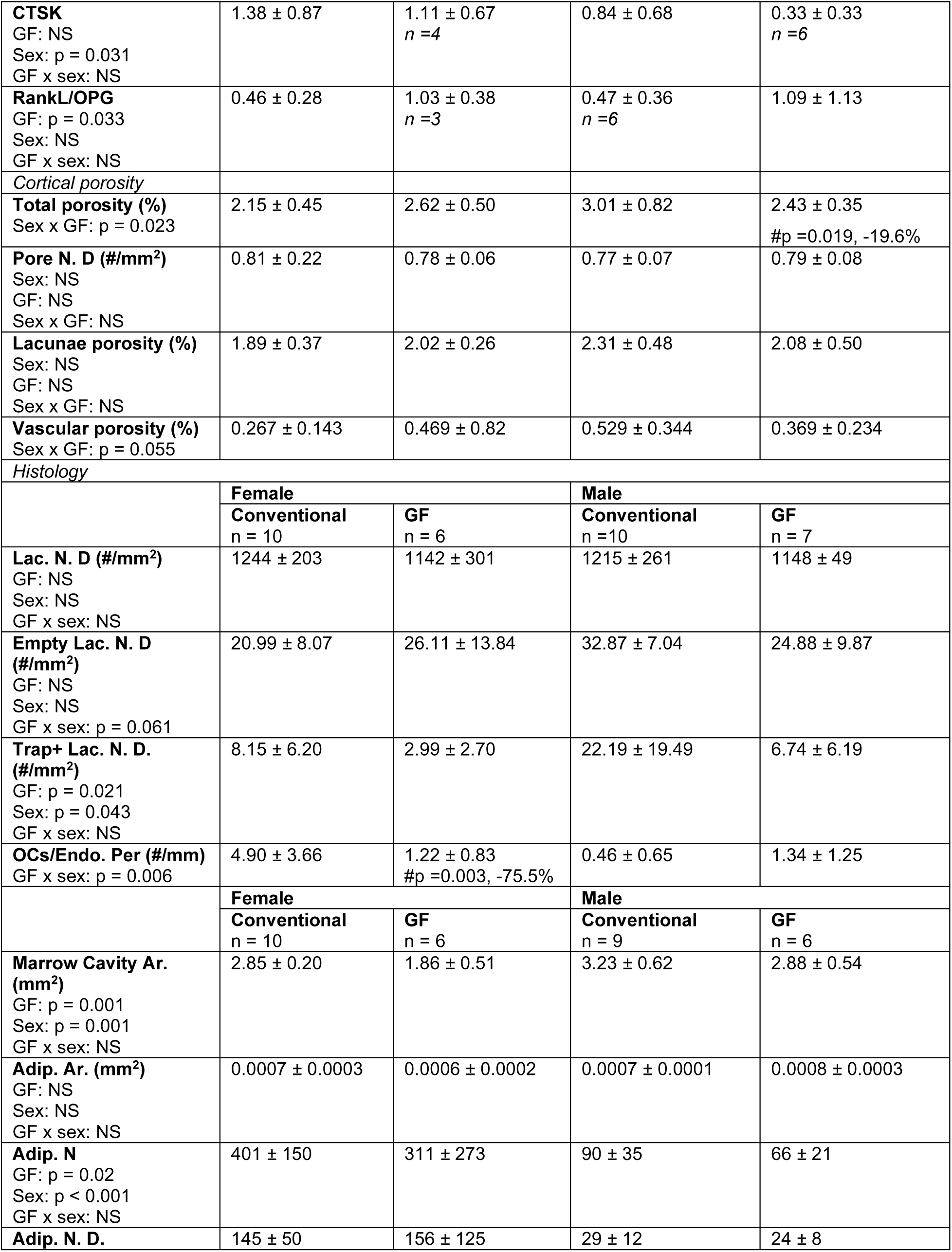
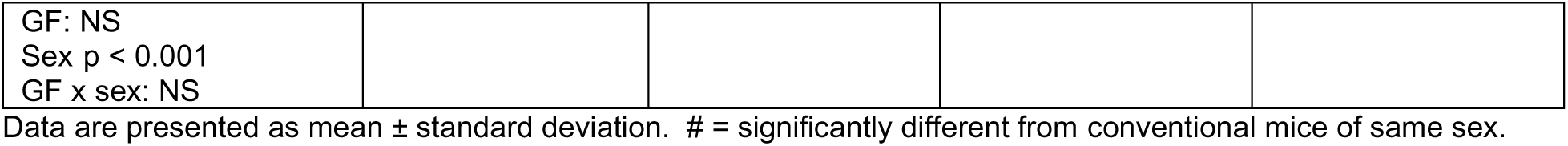
Results from all analyses.

## Supplement 2

Designated for extended metabolomic data.

**Note:** Source mass spectrometry files are available on the Metabolomics Workbench

**Supp. Table 2.**
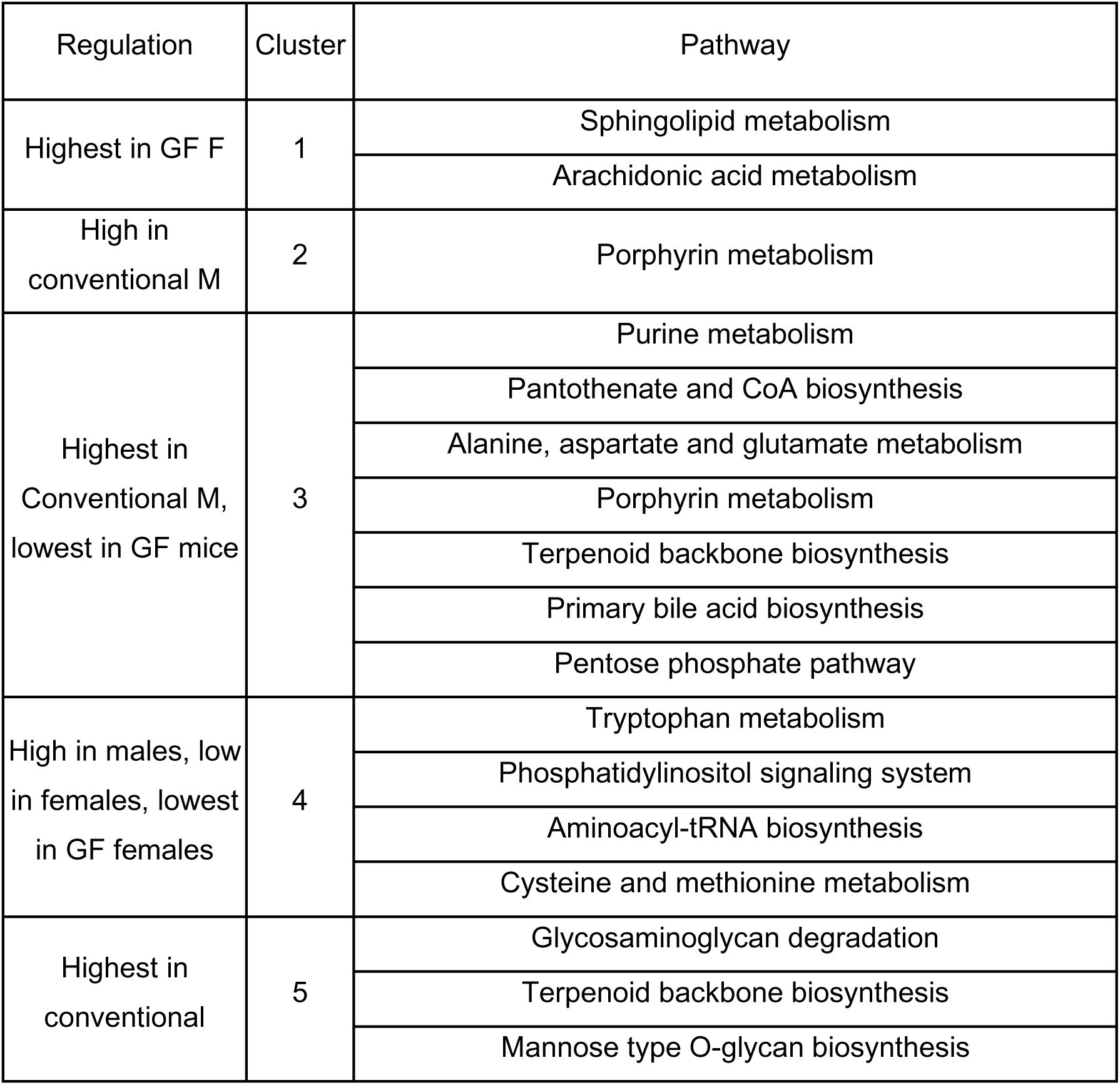
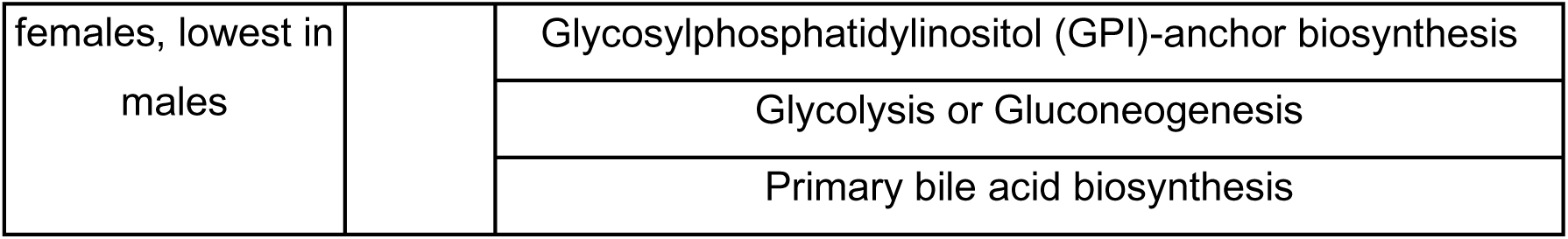
Metabolic pathways associated with GF and conventional male and female mice identified by median metabolite intensity heatmap analysis. Pathways listed have an FDR-corrected significance level < 0.05.

## Notes

### Competing Interest Statement

The authors have declared no competing interest.

